# Early mitochondrial stress and metabolic imbalance lead to photoreceptor cell death in retinal degeneration

**DOI:** 10.1101/2021.10.10.463827

**Authors:** Ke Jiang, Anupam Kumar Mondal, Yogita K. Adlakha, Jessica Gumerson, Angel Aponte, Linn Gieser, Jung-Woong Kim, Alexis Boleda, Matthew J. Brooks, Jacob Nellissery, Donald A. Fox, Robert Balaban, Raul Covian, Anand Swaroop

**Affiliations:** Neurobiology, Neurodegeneration & Repair Laboratory, National Eye Institute, National Institutes of Health, Bethesda, Maryland, USA; Translational Health Science and Technology Institute, National Capital Region Biotech Cluster, Faridabad, India; Proteomics Core Facility, National Heart, Lung and Blood Institute, National Institutes of Health, Bethesda, Maryland, USA; Department of Life Science, College of Natural Sciences, Chung-Ang University, Seoul 156-756, Republic of Korea; Center for Bioinformatics and Computational Biology, University of Maryland, College Park, Maryland, USA; Laboratory of Cardiac Energetics, National Heart, Lung, and Blood Institute, National Institutes of Health, Bethesda, Maryland, USA

**Keywords:** Retinal degeneration, Vision, Phototransduction, *rd1*, Mitochondria, Metabolome, Proteome, Transcriptome, Complex I, Oxidative phosphorylation, Energy homeostasis, Central carbon metabolism, Calcium signaling

## Abstract

Neurodegenerative diseases exhibit extensive genetic heterogeneity and complex etiology with varying onset and severity. To deduce the mechanism leading to retinal degeneration, we adopted a temporal multi-omics approach and examined molecular and cellular events before the onset of photoreceptor cell death in the widely-used *Pde6b^rd1/rd1^* (*rd1*) mouse model. Transcriptome profiling of neonatal and developing rods revealed early downregulation of genes associated with anabolic pathways and energy metabolism. Quantitative proteomics of *rd1* retina showed early changes in calcium signaling and oxidative phosphorylation, with specific partial bypass of complex I electron transfer, which precede the onset of cell death. Concurrently, we detected alterations in central carbon metabolism, including dysregulation of components associated with glycolysis, pentose phosphate and purine biosynthesis. *Ex vivo* assays of oxygen consumption and transmission electron microscopy validated early and progressive mitochondrial stress and abnormalities in mitochondrial structure and function of *rd1* rods. These data uncover mitochondrial over-activation and related metabolic alterations as early determinants of pathology and implicate dysregulation of calcium signaling as the initiator of higher mitochondrial stress, which then transitions to mitochondrial damage and photoreceptor cell death in retinal degeneration. Our studies support the “one hit model” arguing against the cumulative damage hypothesis but suggest that cell death in neurodegenerative disease is initiated by specific rather than a random event.

## INTRODUCTION

Neurodegenerative diseases are characterized by progressive dysfunction and death of post-mitotic neuronal cells that lead to severe cognitive and/or sensory defects, greatly impacting the quality of life. These largely untreatable pathologies typically afflict specific neurons, with divergent clinical manifestations caused by heterogenous genetic factors ^1,2^. Genes and pathways implicated in neurodegeneration participate in energy metabolism, signaling, stress response, and autophagy, among other homeostatic processes ^3–5^. Mitochondrial dysfunction is arguably a key factor in neurodegenerative disease ^6–8^. However, the concept of increased stress of functional mitochondria ^9^ has been largely underappreciated. One carbon and serine metabolism pathways are also implicated in health and pathology and may provide novel therapeutic targets ^10,11^. To date, we still lack knowledge of the precise cellular events, especially integrative molecular cues convergent at the onset of pathology that determine the natural history of neuronal cell death in degenerative disease.

The retina, the most accessible part of the central nervous system, provides an excellent model for examining neurodegeneration. Retinal and macular degenerative diseases constitute a major cause of incurable vision impairment worldwide, with pathogenic mutations identified in almost 300 genes (https://sph.uth.edu/retnet/). Extensive genetic and phenotypic heterogeneity associated with Mendelian retinal diseases ^12,13^ make the design of gene-based therapies particularly challenging ^14,15^. The death of photoreceptor cells, predominantly by apoptosis, is the primary end-point that leads to vision loss in most retinal diseases ^16,17^, and thus anti-apoptotic interventions have been attempted to restore vision ^18^. The “one-hit model” of cell death in neurodegeneration argues against the cumulative damage hypothesis and suggests that mutations create an adaptive state of equilibrium and cell death is initiated by single random event ^19^. We and others hypothesized convergence in pathways leading to pathology in divergent inherited retinal diseases ^20,21^. The prodromal events and critical mutant steady state preceding photoreceptor cell death are poorly understood.

Spontaneous and transgenic mouse mutants can elegantly phenocopy human retinal disease ^22,23^; albeit, the onset and severity may vary. We chose a widely-studied retinal neurodegeneration model, the retinal degeneration 1 mouse (*rd1*; also referred as *Pde6b^rd1/rd1^*), to investigate early molecular and cellular events that may trigger photoreceptor cell death. The *rd1* mouse is homozygous for the loss of function genetic defect in the *Pde6b* gene, which encodes the rod photoreceptor-specific cGMP phosphodiesterase 6 β subunit (PDE6β) ^24,25^. PDE6β is an evolutionarily conserved key component which is localized in rod outer and inner segments ^26^ and mediates phototransduction by controlling cGMP levels and facilitating the opening of cGMP-gated ion channels ^27^. The rod cell death in *rd1* mouse retina is histologically evident starting at postnatal day (P)10, with a complete loss of the photoreceptor outer nuclear layer (ONL: composed of rod and cone nuclei) by P30. Mutations in the human *PDE6B* gene lead to a spectrum of autosomal recessive retinal degeneration phenotypes ^28^.

Here, we report widespread prodromal pathophysiological changes in the neonatal *rd1* retina using an integrative multi-omics approach that included temporal transcriptomics of purified rod photoreceptors along with proteomic and metabolomic analysis of whole retinas in combination with structural and functional assays. These data demonstrate a higher mitochondrial stress as early as P6; much before photoreceptor cell death is evident in the mutant retina. We show, for the first time, that calcium signaling defects which drive mitochondrial and metabolic alterations constitute the pre-death mutant state in rod photoreceptors. These results are consistent with the known high metabolic demand and activity of photoreceptors ^29–31^. Our findings of prodromal molecular and functional dysfunction in mitochondria and their associated cell signaling regulation are the earliest detected etiology of rod cell death and support the proposed convergence of cellular events during retinal neurodegeneration ^2,20,21^.

## RESULTS

### Early transcriptomic divergence in the *rd1* rods reveals altered metabolic and signaling pathways

Gross morphological changes and cell death markers are detected in rod photoreceptors of *rd1* mouse retina starting at or around P10, and early pathology is characterized by shorter outer segments and progressive thinning of the ONL (Figure 1A). Since the genetic and biochemical defect is limited to these retinal cells, we first performed transcriptome profiling of newborn and developing rod photoreceptors. We generated *rd1*-GFP mice by mating the *Pde6b^rd1/rd1^* mice to *TgNrlp-EGFP* (*WT*)^32^ mice, used flow cytometry to purify GFP^+^ rods at P2, P4, P6, P8 and P10 (before rod cell death is evident), and performed RNA-seq analysis (Figure 1A). After rigorous filtering and normalization, we captured expression of 14,534 genes from temporal transcriptome profiles of *rd1* and *WT* photoreceptors. Principal component analysis (PCA) of *rd1* and *WT* samples showed progressively divergent RNA profiles as early as P2 (the peak of rod birth is P0-P2), the earliest time we sampled the purified rods (Figures 1B and 1C). Heat maps of a total of 1,474 differentially expressed genes (DEGs) displayed distinct patterns (shown as clusters C1-C10: see below) in *rd1* rods at pre-degeneration states (Figure 1D).

**Figure 1.**
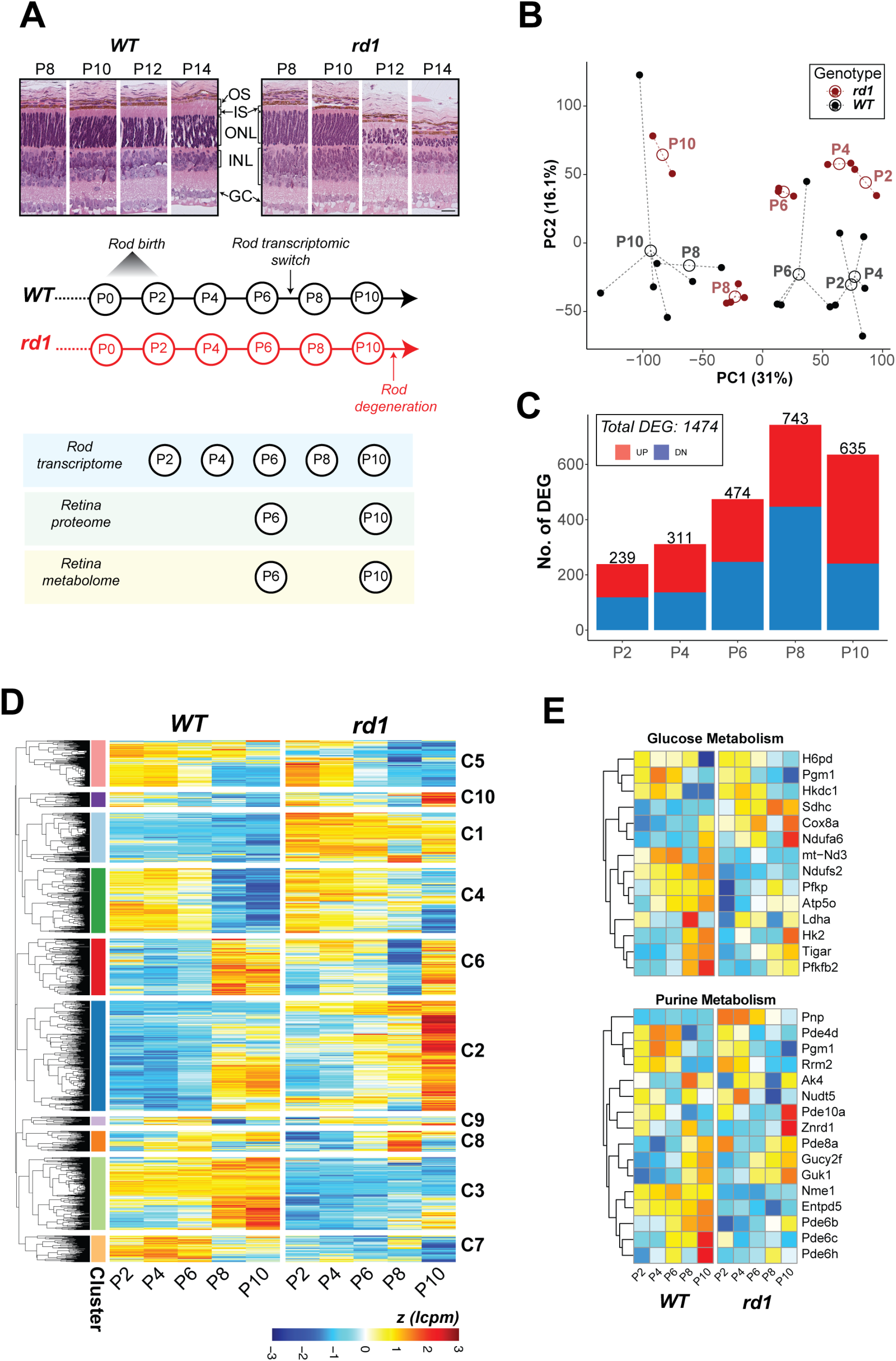
Transcriptome dynamics of rod photoreceptors prior to cell death in the *rd1* retina. A. Pathology in the *rd1* retina and study design. Representative histology pictures of retinal cross sections from *WT* and *rd1* mice at ages ranging from P8 to P14 (Top panel). A schematic summary of timeline showing development in *WT* retina and onset of degeneration in *rd1* (Middle panel). Experimental plan and ages of sampling for each assay (Bottom panel). Scale bar = 20 μm. OS: outer segment, IS: inner segment, ONL: outer nuclear layer, INL: inner nuclear layer, GC: ganglion cell. B. Principal Component Analysis (PCA) plot summarizing the transcriptomic landscape of rod photoreceptors of *WT* and *rd1* retina. C. Trend of differential gene expression between *WT* and *rd1* rods. The number atop each bar represents the total number of significantly differentially expressed genes (DEGs) in a pairwise comparison between age-matched *WT* and *rd1* samples. D. Heatmap of gene expression of all DEGs that are significant across age-matched comparisons. Log CPM (lcpm) values are row scaled to z-scores for plotting. See also Figure S1D. E. Heatmaps of gene expression of significant DEGs that participate in glucose metabolism (glycolysis, oxidative phosphorylation, pentose phosphate) and purine metabolism pathways. Log CPM (lcpm) values are row scaled to z-scores for plotting. Color scale bar is same as in Figure 1D. See also Figure S2.

KEGG pathway enrichment analysis of DEGs revealed that central carbon metabolism and signaling pathways were prominently impacted in the early stages of developing *rd1* rods (Figures 1E, S1 and S2). Metabolism related DEGs at P2, P4 and P6 included those associated with carbohydrate metabolism, cellular anabolic processes, and fatty acid biosynthesis (Figures 1E, S1A-S1C and S2) and corresponded primarily to glycolysis (e.g., *Pfkp, Tigar*, and *Pfkfb2*), oxidative phosphorylation (OXPHOS; e.g., *Ndufs2, Atp5o* and *mt-Nd3*), pentose phosphate pathway (PPP) (e.g., *H6pd*), purine metabolism (e.g., *Pnp*), and glycogenolysis (e.g., *Agl* and *Pgm1*) (Figures 1E, S1B and S2). A number of lipid metabolism genes exhibited higher expression; e.g., fatty acid biosynthesis gene *Acot1* and several phospholipid metabolism genes (*Abhd4, Lpcat2*, and *Samd8*) (Figures S1B and S2). Additionally, homocysteine and one carbon metabolism genes – *Ahcy, Mthfr* and *Mthfd2* – were among those showing the highest fold change differences (Figure S1B, Table SData1). The gene coding for myoferlin (*Myof*) was markedly under-expressed in *rd1* rods (Figure S1B), even though its consequence in neuronal cells remains unspecified. Transcriptional modulators, such as *Duxbl1, Sox30* and *Hmga1b*, and mitochondrial stress regulator *Atf4* were also significantly differentially expressed in the early developing rods (Figure S2). Importantly, we observed differential expression of calcium-related signaling genes (e.g., *Hcn, S100a10, Atf4*) in *rd1* rods at early stages, much before the onset of degeneration at P10 (Figures S1C and S2). Finally, genes involved in the phototransduction pathway demonstrated significantly altered expression at these early ages (Figure S1A).

*Rd1* rod DEGs could be grouped into 10 clusters (C1-C10). Three of the larger clusters showed concordance with changes in distinct pathways (Figures 1D, S1D and S1E). Cluster 1 (C1) genes consistently demonstrated higher expression in *rd1* rods starting at P2 and included metabolic genes for one carbon pool, purine metabolism, and fatty acid metabolism. Conversely, genes in C3 cluster with low expression in *rd1* rods belonged to phototransduction and hedgehog signaling. Cluster C2 showed sharp induction of gene expression at the onset of degeneration (P10) and contained fatty acid biosynthesis, ferroptosis as well as phototransduction genes.

### Reduced expression of mitochondrial NADH dehydrogenase complex and signaling proteins in *rd1* retina

In developing mouse retina, the photoreceptor differentiation is concordant with major changes in the transcriptome between P6 and P10 ^33^. We therefore wanted to ascertain whether disease-specific trends in *rd1* gene expression further translate to proteome changes that precede developmental milestones and onset of degeneration. Mass spectrometry-based proteome analysis of P6 and P10 *rd1* and *WT* retina identified a total of 4,955 proteins. Of these, 621 were differentially expressed in P6 *rd1* retina and an additional 630 showed altered expression at P10 (Figures 2A and S3A). Gene set enrichment analysis (GSEA) using Fast Gene Set Enrichment Analysis (*<u>fgsea</u>*) uncovered mitochondrial electron transport chain (ETC) as the most downregulated pathway in both P6 and P10 *rd1* retina (Figures 2B and S3B) with specific reductions in the subunits of NADH dehydrogenase (complex I) (Figure 2C). Further targeted analysis, using a curated GSEA, confirmed a significant negative enrichment of the OXPHOS pathway at P6 and P10 (Figure S3C). This trend begins a negative course at P6 with further reductions by P10, clearly showing a correlation of OXPHOS impairment with age in *rd1* retina. Leading-edge analysis identified complex I subunits to be the driver of OXPHOS impairment, with other ETC complexes showing insignificant changes (Figure 2D). Complex I enzymatic activity assay in mitochondria-enriched retinal extracts also showed lower activity in *rd1* retina between P2 and P8 (Figure S3D), confirming an impairment of complex I function.

**Figure 2.**
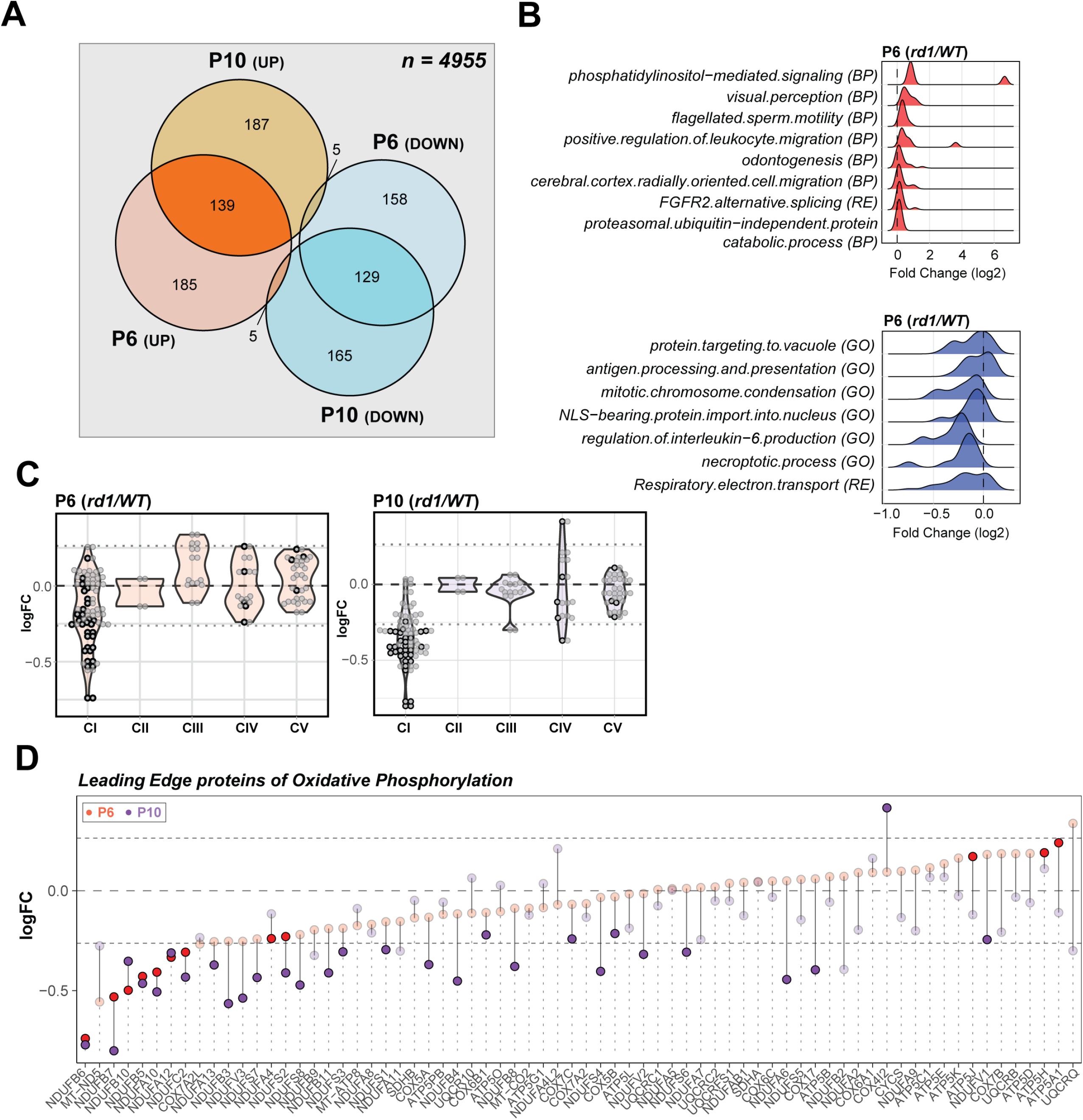
Proteomics identifies dysregulation of Electron Transport Chain complex I in the *rd1* retina before degeneration onset. A. Venn diagram comparison of differentially expressed proteins in P6 and P10 *rd1* retina. B. Summary of pathway enrichment analysis that identifies differential regulation at early stages (P6) in the *rd1* retina. Ridge-plots show fold change (*rd1* vs *WT*) distribution of leading-edge proteins from significant genesets. The top and bottom panels show over- and under-enriched pathways in red and blue colors respectively. See also Figure S3B for late stage (P10) data. C. Proteomic fold change (FC) of mitochondrial electron transport chain complexes. Black dots inside violin plots represent significant differential expression of an individual protein (*rd1* vs *WT*) of a specific complex. Gray dots inside violin plots represent non-significant differential expression. D. Dumb bell plot showing change in protein expression from P6 to P10 for leading-edge proteins of oxidative phosphorylation. Dark red and purple dots represent significantly different proteins at P6 and P10, respectively. Grayed dots represent proteins with expression less than statistical significance threshold. See also Figure S3C.

Manual curation of significantly altered mitochondrial proteins uncovered overexpression of several ribosomal proteins, tRNA ligases, import/processing subunits and ubiquinone biosynthesis proteins at P6 and P10, suggesting a more active mitochondrial biogenesis in *rd1* retina (Figure 3A). For example, the most over-expressed mitochondrial protein (SUPV3L1, FC_P6_ = 4.98) is an RNA helicase associated with the RNA surveillance system in mitochondria ^34^, suggesting enhanced transcription of mitochondrial DNA. In addition, several proteins involved in assembly of respiratory complexes were increased in *rd1* retina (ATPAF1, TTC19, AIFM1 and NLN). Furthermore, increased expression of amino acid degradation enzymes (GLDC, IVD, MCCC2) suggested an augmented degradation of mitochondria in *rd1* retina, resulting in a faster mitochondrial turnover due to a parallel increase in biogenesis. In addition, mitochondria in the mutant retina seem to exhibit a higher capacity to donate electrons directly into the ubiquinone pool bypassing complex I, as suggested by higher expression of enzymes involved in fatty acid oxidation (MCEE, ECI2, HADHA, ECH1, ECI1), leucine degradation (IVD, MCCC2) and pyrimidine metabolism (DHODH).

**Figure 3.**
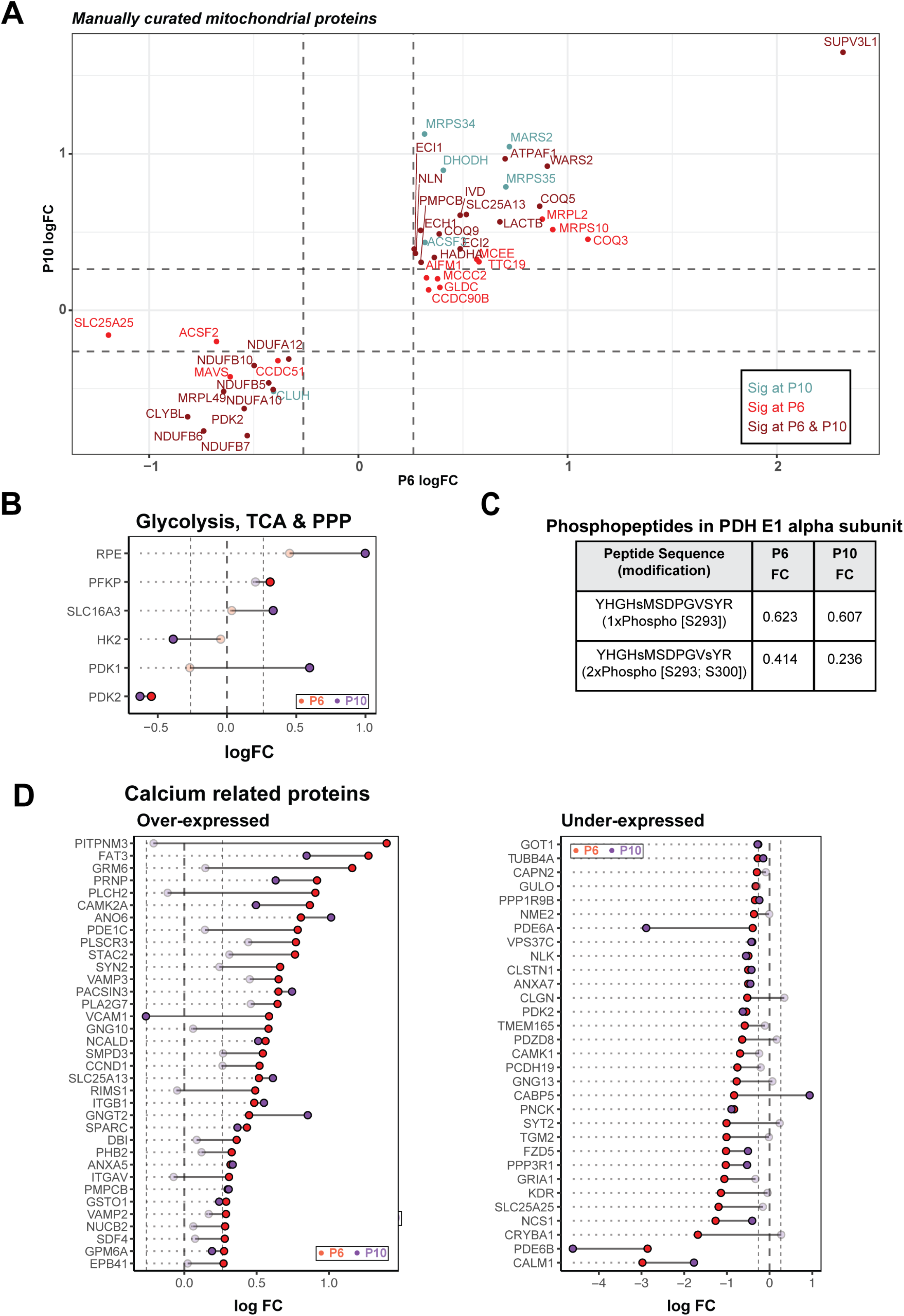
*rd1* retinas harbor abnormal abundance of mitochondrial and calcium related proteins before degeneration. A. Scatter plot of manually curated mitochondrial proteins show differential regulation of mitochondrial metabolic and signaling proteins at P6 and P10. Proteins with abundances significantly different at P6, P10, and both P6 and P10 are labeled in red, blue, and brown, respectively. B. Dumb bell plot showing change in protein expression from P6 to P10 for significantly differential glycolysis, TCA and PPP proteins. Dark red and purple dots denote proteins above significance threshold at P6 and P10, respectively. Gray dots represent proteins with non-significant differential expression. C. Phosphopeptides detected in pyruvate dehydrogenase (PDH) E1 alpha subunit. D. Dumb bell plot showing change in protein expression from P6 to P10 for significantly differential calcium related proteins. Plot on the left summarizes proteins overexpressed in the *rd1* retina, whereas plot on the right depicts under-expressed proteins. Color codes are same as in Figure 3B.

Curiously, a similar analysis of the strategically upstream glycolytic, PPP and TCA cycle proteins did not show a pathway-wide consistent difference, although a few specific proteins including PFKP, SLC16A3, HK2, PDK1 and PDK2 were significantly different at one or both ages (Figure 3B). Consistent with the significant under-expression of the pyruvate dehydrogenase kinase PDK2 at P6 and P10, we observed reduction in ratios of single- and double-phosphorylated peptides from subunit E1α of the pyruvate dehydrogenase complex (PDC) in *rd1* at both P6 and P10 (Figure 3C). When analyzing the differential protein expression of all calcium-associated proteins, robust and consistent calcium dysregulation was evident in the *rd1* retina (Figure 3D). One of the most downregulated proteins in the *rd1* retina was calmodulin (CALM1) (Figure S3A), which serves as the primary intracellular calcium sensor and initiator of downstream signaling events that correspond to changing calcium levels. In addition, calcium-binding mitochondrial carrier proteins, such as SLC25A13 (fold change at P6, FC_P6_ = 1.48) and SLC25A25 (FC_P6_ = 0.43), were significantly altered. Interestingly, the highest overexpressed protein in *rd1* retina was PITPNM3 (FC_P6_ = 2.63), a membrane-associated phosphatidylinositol transfer protein implicated in autosomal dominant cone rod dystrophy ^35^. These findings strongly indicate abnormal calcium signaling in the *rd1* retina at early stages before the onset of degeneration.

### Metabolite imbalance especially of central carbon pathways in *rd1* retina

To further dissect early stages of neurodegeneration, we examined 116 metabolites in *rd1* and *WT* retina at P6 (Figure 4) and P10 (Figure S4). We detected alterations in glycolytic and PPP intermediates as well as nucleotides at both stages (Figures 4A and S4A). Carnosine, a dipeptide involved in resistance to oxidative stress and divalent ion chelation ^36^ was the most elevated metabolite in the P6 *rd1* retina. We generated a metabolite-pathway network and identified biosynthesis of amino acids, carbon metabolism, ATP-binding-cassette transporters, purine metabolism and PPP as those with the most difference in metabolites (Figure 4B).

**Figure 4.**
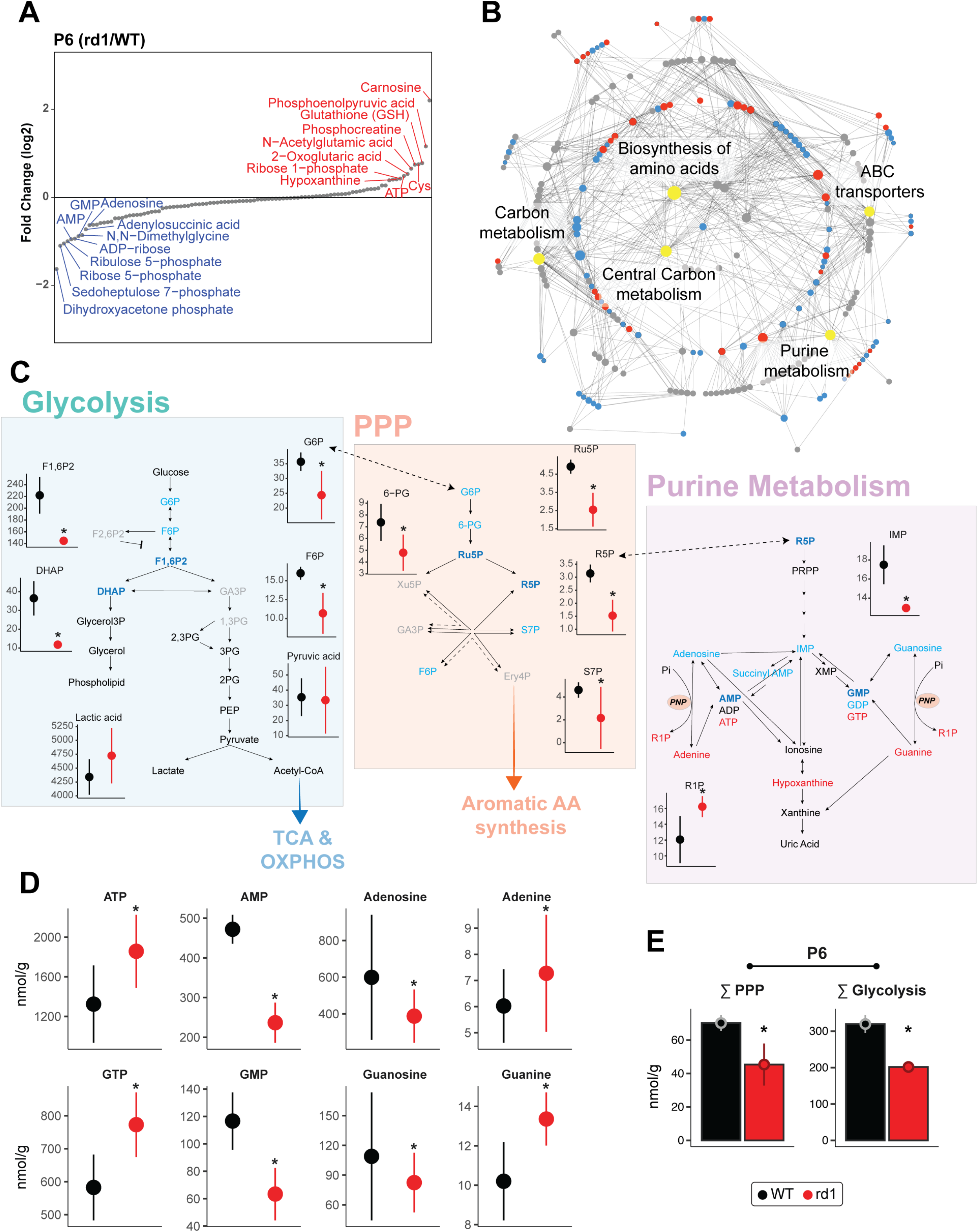
Global metabolite profiling identifies aberrant central carbon metabolism. A. Relative abundance plot for the metabolome of the *rd1* retina relative to *WT*, at P6, showing the highest and least abundant metabolites. See also Figure S4A for P10 data. B. Mapping of identified metabolites to KEGG pathways. Significantly altered metabolites at P6 in *rd1* relative to *WT* retinas are colored in either red or blue corresponding to increased or decreased abundance, respectively, while pathways are shown in grey. The top five pathways with the most significantly different metabolites are labeled and highlighted in yellow. Grey dots are metabolites with differential abundance level less than statistical significance threshold. C. Annotated pathway diagram and connections between glycolysis, PPP, and purine metabolism, with differential metabolites at P6 highlighted. Metabolites with significantly different abundance (*p*<0.05) are highlighted in dark blue, while those showing only differential abundance higher than fold change threshold (>1.2 fold) are highlighted in light blue. Black colored metabolites are detected but do not show differential abundance, while gray colored metabolites were not detected in the metabolomic analysis. Red and black colors in metabolite abundance plots represent *rd1* and *WT*, respectively. Significantly altered metabolites are marked with an asterisk (see STAR Methods for details). D. Abundance levels of adenine and guanine and their corresponding nucleotides at P6. Color and significance features are same as Figure 4C. E. Sum of metabolites involved in glycolysis and PPP in the *rd1* retina at P6. Color and significance features are same as Figure 4C. See also Figure S4B for P10 data.

We then focused on central carbon metabolism (i.e., glycolysis, PPP and purine metabolism) because of significant aberrations in the P6 *rd1* retina. We observed a significant decrease in the upstream intermediates (glucose 6-phosphate, fructose 6-phosphate, fructose 1,6-biphosphate and dihydroxyacetone phosphate) (Figure 4C), yet downstream glycolytic intermediates and products such as glycerol 3-P and glycerol (required for phospholipid synthesis) as well as pyruvate and lactate were relatively unchanged. Together, these data suggest that the net glycolytic flux did not decrease at this early stage (i.e., by P6). Our results also demonstrated significant depletion of most PPP intermediates in the *rd1* retina. In contrast, we observed a marked increase of purine triphosphates and their respective nucleobases, apparently at the expense of early purine biosynthesis intermediates including adenosine and guanosine and their monophosphorylated forms (Figure 4D). Thus, significant reductions of total metabolite concentrations in glycolysis and PPP were observed in the *rd1* retina as early as P6 and even at P10 (Figures 4E and S4B). These metabolic changes suggest a shift in carbon utilization rather than a decrease in metabolic flux through these pathways.

### Pre-degeneration functional and morphological defects in *rd1* mitochondria

Higher ATP concentrations in P6 *rd1* retina (see Figure 4D) suggested higher OXPHOS activity. To directly assess mitochondrial respiration coupled to ATP production in relation to maximal (uncoupled) respiratory activity, we measured oxygen consumption rate (OCR) in acutely isolated, *ex vivo* retinal punches from *rd1* and *WT* mice (Figure 5A) using a previously optimized protocol ^9^. We noted that basal mitochondrial OCR (coupled to ATP production) decreased in *WT* retina during development, reaching a low point at P10, followed by an increase at eye opening on P14 (Figure 5B). In contrast, the *rd1* retinas showed a trend of higher basal OCR compared to *WT* as early as P3 (Figure 5B). With the onset of degeneration after P10 (see Figure 1A), basal mitochondrial OCR in *rd1* retina was reduced significantly at P14 (Figure 5B) when as many as 50% of the rod photoreceptors are lost. After adding a mitochondrial uncoupler (Bam15) to obtain the maximal OCR, the mitochondrial reserve capacity (MRC) was obtained by calculating the additional percentage of maximal OCR that is not used for ATP production under basal conditions. In developing *WT* retina, MRC progressively increased and peaked to ~50% around P8 and P10 before decreasing to ~30% at P14 (Figure 5B), consistent with our previous observations of low MRC in healthy adult retina ^9^. Interestingly, *rd1* retinas were characterized by lower MRC values from P3 through P10 before overt rod degeneration was observed at P14. MRC was markedly increased at P14 by 50-60%, as previously reported ^9^ (Figure 5B).

**Figure 5.**
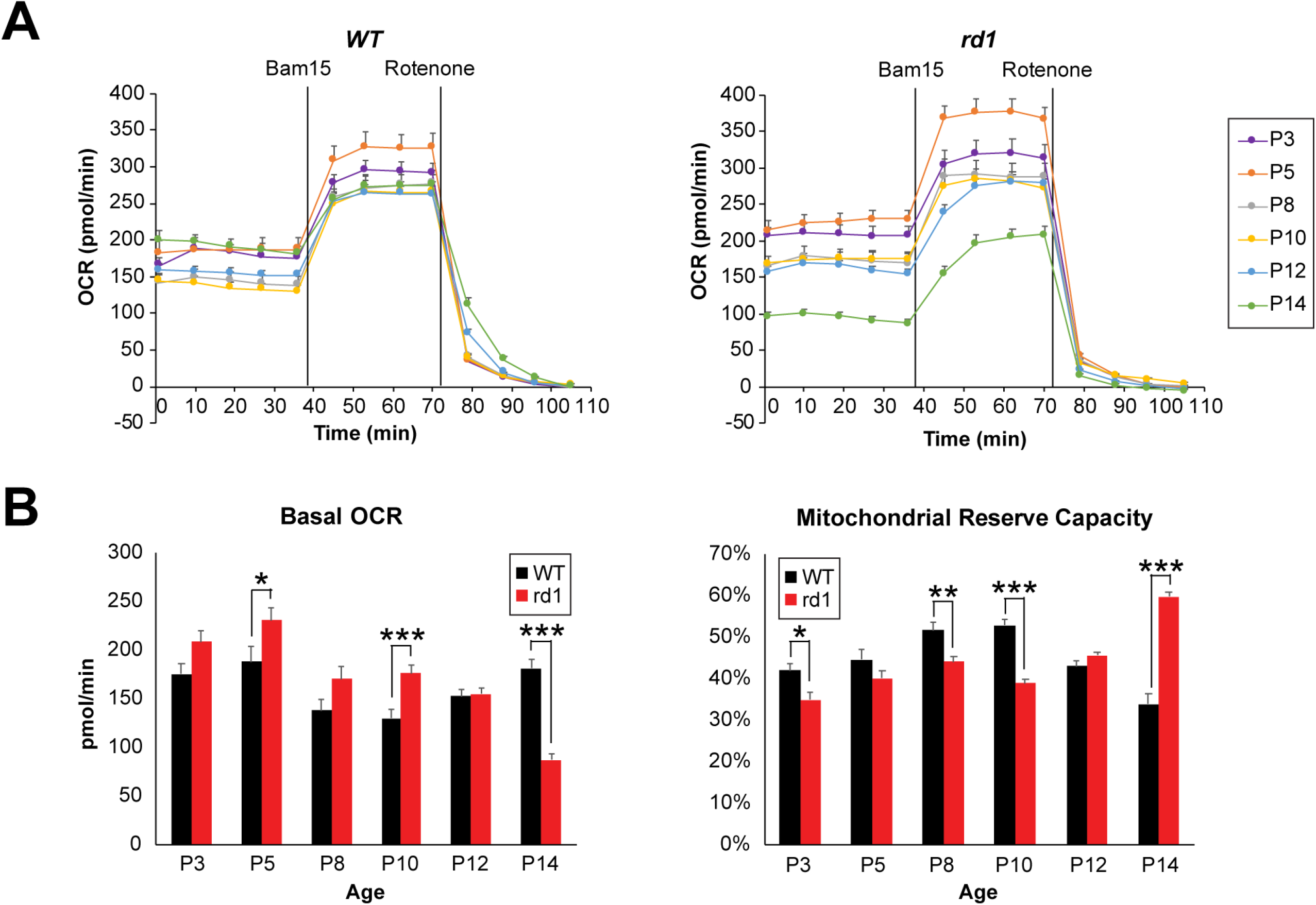
Early functional aberrations in mitochondria before the onset of photoreceptor degeneration in *rd1* retina. A. Traces of mitochondrial OCR in retinal punches isolated from *WT* and *rd1* mice at various ages from P3 to P14 (n=12 to 29 retinal punches, for each group). BAM15 and rotenone were added at indicated time during experiments. Data are represented as mean ± SEM. B. Basal mitochondrial OCR in *WT* and *rd1* retinal punches at each age tested. Values were taken at 36 min, right before addition of BAM15. Plot of mitochondrial reserve capacity in *WT* and *rd1* retinal punches at each age tested. Data are represented as mean ± SEM. Student t test, * *p*<0.05, ** *p*<0.01, *** *p*<0.001.

To assess whether enhanced oxygen consumption might result from increased number of mitochondria in the *rd1* retina, we examined the Mit/Nu DNA ratio ^37^. Quantitative qPCR analysis demonstrated a trend of lower Mit/Nu DNA ratios in *rd1* retina, suggesting relatively fewer mitochondria in P2 to P8 *rd1* retina compared to age-matched *WT* retina (Figure S5).

Ultrastructural analysis by transmission electron microscopy (TEM) revealed delayed organization of the rod inner segments and morphological defects in mitochondria cristae structure as early as P6 in the *rd1* retina (Figure 6). At P8, abnormal mitochondria were more prevalent in inner segments of *rd1* rod photoreceptors and showed loss of inner membrane cristae and a swollen vesicular appearance. However, a majority of mitochondria at this stage still exhibited normal gross morphology. Swollen mitochondria were larger and more numerous in *rd1* rods at P10, indicating a disrupted mitochondrial network even before the histological rod cell death was observed. With extensive degeneration evident at P14, few mitochondria in *rd1* rods had a normal gross or ultrastructural morphology, and the retina exhibited a disrupted overall structure and lamination (Figure S6).

**Figure 6.**
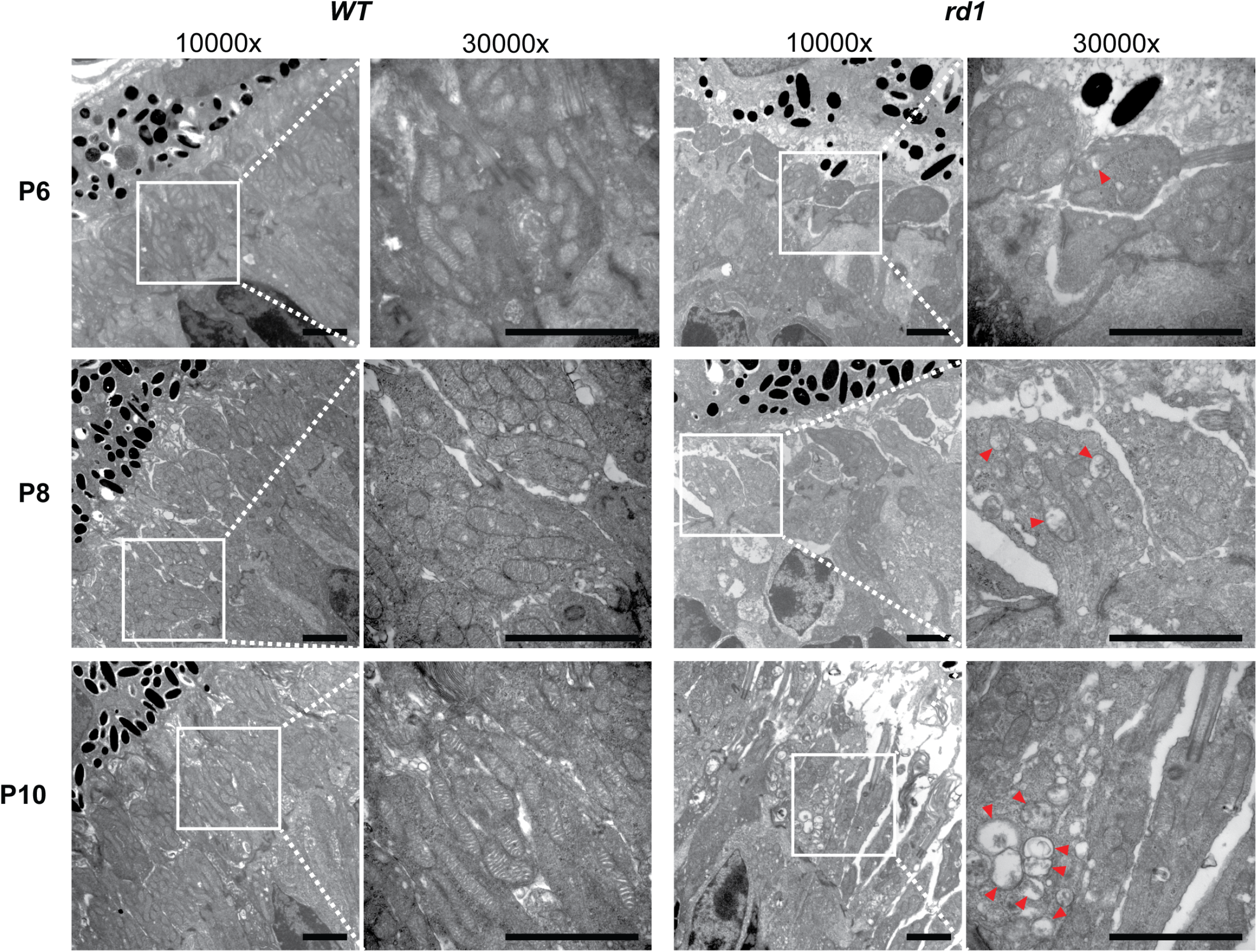
Mitochondrial ultrastructural alterations at early stages before photoreceptor degeneration in *rd1* retinas. TEM of photoreceptor inner segment area from P6 to P10 *WT* and *rd1* retina are shown at magnifications of 10,000x and 30,000x (zoomed-in area, indicated by white square box). Red arrowheads indicate abnormal mitochondria with swollen cristae. Scale bar = 2 μm. See also Figure S6.

### Proposed model of molecular and cellular events leading to photoreceptor cell death

Temporal dynamics of transcriptome changes, together with proteomic and energy metabolism defects detected at early stages in the *rd1* retina, enabled us to construct a model to explain the etiology of rod photoreceptor cell death in the *rd1* mouse (Figure 7). Important features identified in our study include progressively increasing metabolic alterations and mitochondrial deficiencies that were initiated prior to any observed morphological and/or functional defects in degenerating rod photoreceptors of the *rd1* mouse. We hypothesize that a functional imbalance in pathways associated with one-carbon metabolism and energy homeostasis, as indicated by RNA profiles of newborn rod photoreceptors, results in down regulation of complex I, and aberrant mitochondrial structure and function. As a consequence of reduced electron flow through the ETC from NADH oxidation, the ubiquinone pool receives more electrons from other sources such as fatty acid oxidation, as well as glycine, leucine and valine metabolism. Concurrently, expression levels of a key intracellular calcium regulator, calmodulin, are greatly reduced leading to dysregulation of mitochondrial calcium uptake and calcium-dependent signaling pathways (e.g., the activation of PDC through calcium-activated dephosphorylation by PDP). Activated PDC draws from the pyruvate pool and can enhance the flux through glycolysis pathway. A decrease in upstream metabolites of both glycolysis and PPP supports this hypothesis. Furthermore, disease-related activity of the one-carbon cycle and homocysteine metabolism can aggravate cellular distress by potentially altering NADPH levels, consistent with observed differences in purine metabolism. In parallel, later accumulation of cGMP in *rd1* mutants may also lead to differential abundance of purine molecules, reflecting the rod cell’s attempts to control purine production by metabolic reorganization, which can further contribute to initiation of molecular pathology. Thus, an increasing and accumulating imbalance of energy metabolism pathways coupled with a progressive increase in intracellular and mitochondrial calcium levels result in elevated mitochondria stress and calcium overload, eventually leading to mitochondria swelling and rod photoreceptor cell death in the *rd1* retina.

**Figure 7.**
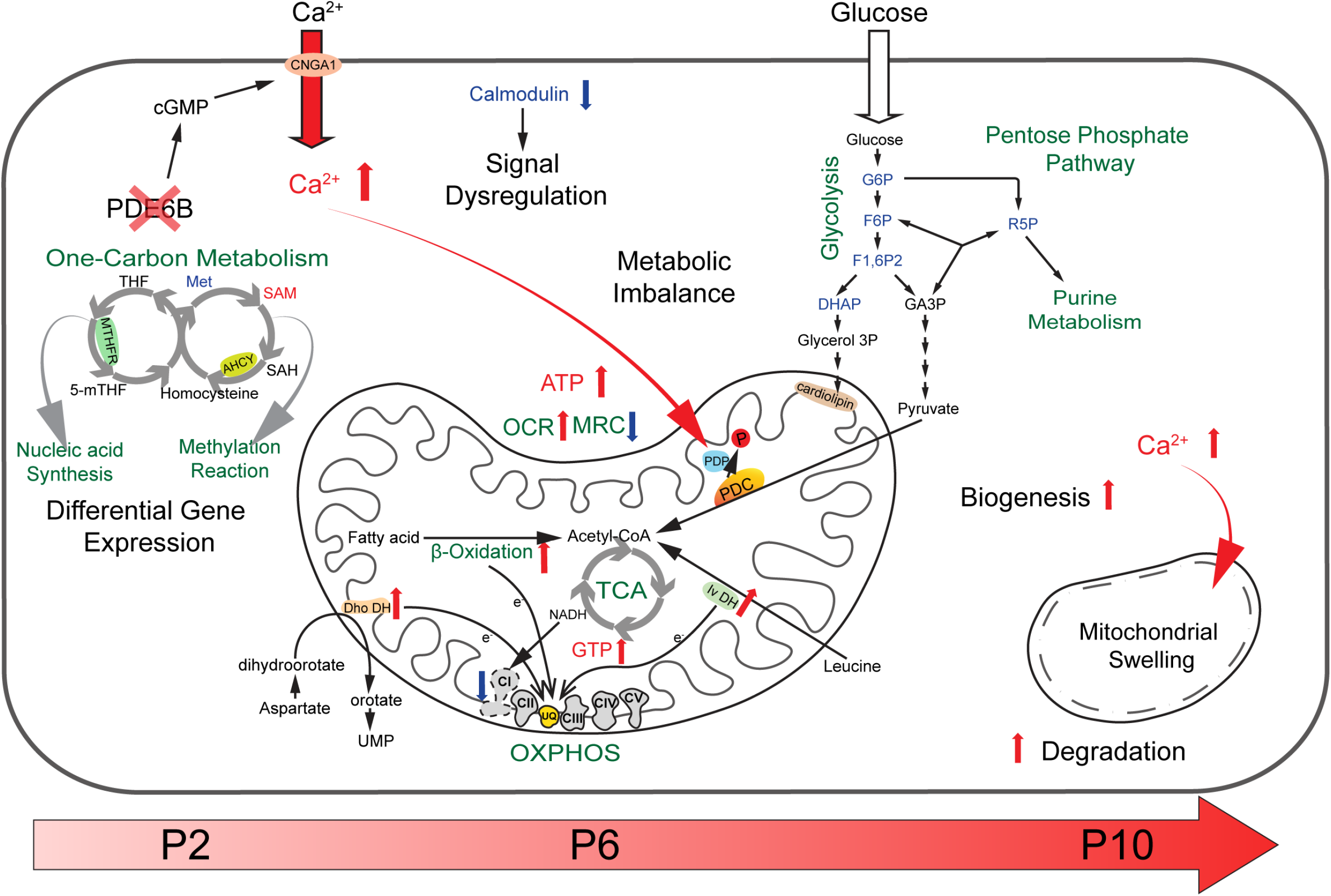
An integrated mitochondria-centered model of molecular events leading to photoreceptor cell death in retinal degeneration. Altered gene expression starts in newborn *rd1* rod photoreceptors carrying the *Pde6b* mutation. Differentially expressed key enzymes (MTHFR and AHCY) affect the balance of one-carbon metabolism, which in turn influences nucleic acid synthesis and pan-methylation status. Concurrently, reduced calmodulin expression leads to dysregulation of calcium-mediated signaling. Increased intracellular calcium enters mitochondria, activates PDP, which in turn stimulates PDC by enhancing its dephosphorylation. Faster consumption of pyruvate triggers an imbalance of the metabolite pool in glycolysis and pentose pathways. Higher calcium also increases damage to mitochondria, which are replaced more rapidly by accelerated degradation and biogenesis (turnover). Enhanced biosynthesis of mitochondrial membrane phospholipids would further deplete glycolytic intermediates. Lower complex I activity is compensated by increased electron transfer from fatty acid, aspartate and leucine metabolism into the UQ pool. These changes lead to enhanced OCR, reduction in MRC, and increase in cytosolic ATP. With progressive imbalance in metabolism and augmented stress, mitochondria and their cristae swell and rupture, eventually committing the cell to a death fate. UQ: ubiquinone, Dho DH: dihydroorotate dehydrogenase, Iv DH: isovaleryl-CoA dehydrogenase, PDC: pyruvate dehydrogenase complex, PDP: pyruvate dehydrogenase phosphatase, PDK: pyruvate dehydrogenase kinase.

## DISCUSSION

The lack of regenerative potential in the mammalian retina makes photoreceptor cell dysfunction and/or death a leading cause of incurable blindness. Rod photoreceptors are specifically vulnerable to degeneration, and their deficiency is generally followed by the loss of cones in inherited retinal diseases ^21^. Even in a multifactorial disorder affecting the macula (namely age-related macular degeneration, AMD), rod cell death precedes cone loss ^38^. Convergence to rod degeneration despite extensive clinical and genetic heterogeneity suggests commonalities in molecular and cellular pathways and offers opportunities for gene-agnostic intervention in retinal and macular diseases ^20,21^. As an example, a thioredoxin-like protein RdCVF has been shown to protect cone function in multiple distinct models of retinal degeneration by inducing glucose uptake and aerobic glycolysis ^39^. We hypothesized that the presence of a genetic mutation would exert specific impact(s) on molecular/biochemical pathways in target cells early in development and that progressive and varying deterioration of cellular responses would eventually lead to cell death ^20^. By applying three ‘omics’ technologies, here we demonstrate that the loss of PDE6β by an inherited mutation results in extensive adaptations in metabolic and mitochondrial components of newborn and developing rod photoreceptors, much before the established critical requirement of PDE6β in signaling during phototransduction. Our studies identify early convergence of multiple cellular responses to mitochondria-related metabolic and signaling pathways and point to disruptions or reallocations of energy resources and calcium signaling in early stages preceding the onset of retinal neurodegeneration.

The *rd1* mouse represents arguably the most studied model of retinal degeneration. PDE6β is associated with controlling cGMP levels, and cGMP accumulation has been implicated in driving the death of rod photoreceptors in the *rd1* retina as well as in phototransduction-associated inherited retinal diseases ^17,40,41^. cGMP is a key second messenger in multiple cellular signaling pathways. In photoreceptors, cGMP-gated channels control the flux of ions in light and dark conditions contributing to Ca^2+^ homeostasis ^42^. Remarkably however, photoreceptor cell death begins in *rd1* retina around P10 and before eye opening in mouse (at P14). Whilst most studies focused on the disease progression at or after onset of rod death (after P10), our transcriptome profiling of neonatal (P2-P6) *rd1* rods shows major and highly specific abnormalities in expression of genes associated with central carbon metabolism. These early changes likely impact energy homeostasis, redistribute metabolites within one-carbon pathways, and initiate a cascade of signaling events. Two of the earliest differentially expressed genes – *Ahcy* (lower expression) and *Mthfr* (high expression) – regulating the one-carbon metabolism are significantly altered with high magnitude and influence nucleic acid synthesis and the balance of S-adenosylmethionine to S-adenosylhomocysteine (SAM/SAH) ratio. Continued differential expression of *Ahcy* and *Mthfr* is reflected by an increase in SAM in the metabolome data from P10 *rd1* retina, suggesting a shift in pan-methylation status. The differential gene expression trends suggest that mutant photoreceptors never undergo normal development despite their normal morphological appearance in early stages. Proteomic and metabolomic data from P6 *rd1* retina further supports these findings and implicates mitochondria and related metabolic and calcium signaling pathways as the early mediators of the cellular response in retinal neurodegeneration.

Photoreceptors have high energy requirements for maintaining their physiological state in dark and light ^43^. Like other neurons, glucose provides the primary source of energy in photoreceptors ^9,44^, and the stimulation of glucose uptake and/or metabolic intermediates can be neuroprotective in photoreceptor degeneration ^39,45–47^. Specialized cristae architecture, low reserve capacity and high OXPHOS of mitochondria ^9,44,48^ validate their primary role in providing the energy requirements of photoreceptors. Distinctive reduction in complex I subunits and alterations in several mitochondrial proteins (including others associated with ETC complexes) in P6 *rd1* retina strongly argue in favor of altered mitochondrial structure and function being early indicators of photoreceptor stress. Predictably, we detected lower complex I activity in *rd1* retina, consistent with a previous report ^49^. However, a partial deficiency in complex I activity can be compensated by alternate pathways that donate electrons at the level of the ubiquinone pool. Several of these pathways are augmented in *rd1* retina as indicated by increase in proteins associated with fatty acid oxidation (MCEE, ECI2, HADHA, ECH1, ECI1), leucine degradation (IVD, MCCC2) and pyrimidine metabolism (DHODH). These results of partial bypassing of complex I electron transfer are in accordance with the higher basal mitochondrial OCR used for ATP production and enhanced ATP content.

As an indicative parameter of mitochondrial health/stress, the MRC can be affected by several factors, including the ability of the cell to deliver substrate(s) to mitochondria and functional capacity of enzymes involved in electron transport ^50^. Reduced MRC in *rd1* retina as early as P3 suggests that a fraction of mitochondria in mutant photoreceptor cells have undergone uncoupling, as reported in cells during injury and aging as an adaptive response to decrease reactive oxygen species (ROS) ^51^. Changes in MRC are also correlated with oxidative stress, calcium overload and cone cell death ^52^. A lower MRC and higher OCR in the *rd1* retina are consistent with chronically higher calcium concentrations in photoreceptors, as reported as early as P5 ^53^, which would at first stimulate OXPHOS ^54^ and ATP production but then cause swelling and mitochondrial damage by increasing ROS formation and inner membrane depolarization. Moreover, a faster flow of electrons to the ubiquinone pool to compensate for partial complex I deficiency may also result in enhanced ROS. Our ultrastructural observations show progressive mitochondrial damage in developing *rd1* rods. The higher content of proteins involved in biogenesis and degradation suggests a faster turnover to regenerate functional mitochondria. The observed lower Mit/Nu DNA ratios in *rd1* retina supported this hypothesis. With progressive calcium overload, mitochondrial respiration is inhibited and accompanied by cytochrome c release, decreased ATP production and disruption of the mitochondrial matrix and cristae ^55^.

High aerobic glycolysis is a characteristic feature of photoreceptor cells and crucial for anabolic demands and functional homeostasis ^30,56^. In our study, glycolytic abnormalities in pre-degeneration stage *rd1* photoreceptors are noted by significant over-expression of two important regulators: *Tigar* and *Pfkfb2*. TIGAR is a negative regulator of glycolysis, whereas PFKFB2 modulates the pathway by controlling the pool of fructose-2,6-bisphosphate ^57^. Though we did not detect significant change in pyruvate, lactate or acetyl-CoA, the evidence from proteomic profiling indicates re-programming of the Tricarboxylic Acid (TCA) cycle in the *rd1* photoreceptors. Mitochondrial PDC links glycolysis and the TCA cycle, modulating the overall rate of oxidation of carbohydrate fuels under aerobic conditions ^58^. Significantly lower content of one isoform of PDC kinase (PDK2) and reduced phosphorylation of the PDC E1 alpha subunit in P6 and P10 *rd1* retina suggest a faster-working PDC, which can potentially accelerate flux through the TCA cycle. Our data are concordant with previously reported prolonged photoreceptor survival in mouse models of retinal degeneration by increasing glycolysis flux through inhibition of SIRT6 ^46^ or by promoting lactate catabolism as fuel by Txnip administration ^47^. We propose that the faster PDC activity, besides being the consequence of higher mitochondrial calcium, could be an adaptive response of mutant photoreceptors by fine tuning TCA cycle in dealing with the stress. A recent report demonstrating slowdown of disease progression in *Pde6α* and *Rho^P23H^* mouse models by dietary supplementation of TCA cycle intermediates, α-ketoglutarate and citrate, also supports our hypothesis ^59^.

Elevated intracellular calcium cannot be explained by PDE6β’s role in phototransduction at early stages before eye opening. Notably, our metabolomic data does not reveal a significant difference in cGMP concentration between P6 or P10 *WT* and *rd1* retina, even though mitochondrial structural defects were observed as early as P6 and P8. Interestingly, calmodulin, a key regulator of numerous calcium-sensitive proteins, is nearly undetectable at P6 in the *rd1* retina, suggesting a dysregulation of calcium-dependent signaling at an early stage. Given that calmodulin directly interacts with PDE ^60^, we propose that the loss of PDE6β compromises calmodulin stability or activity, and in turn, affects other calcium-dependent processes. Indeed, our proteomic data reveals altered expression of a number of calcium-regulated proteins in the *rd1* retina. Furthermore, calmodulin controls the sensitivity of rod cGMP channels to cGMP in a calcium-dependent manner, and with a decrease in calmodulin, cGMP channels exhibit a higher affinity for cGMP and more readily open to allow the influx of calcium into the inner segments ^61^. This is a more feasible explanation for the initiation of calcium-mediated mitochondria injury that we detect as early as P6. The absence of PDE6β activity during phototransduction likely amplifies the damage at eye opening (P14), resulting in rapid photoreceptor loss in the *rd1* retina.

Our studies argue against the cumulative damage and support an adaptive mutant state in concordance with the “one-hit” model ^19^. Identification of early disease-associated trends reported herein establishes mitochondria as an integrative node in the cellular response to genetic mutations with a potentially decisive role before the onset of neurodegeneration. We surmise that a mitochondrial link to *rd1* specific PDE6β mutation, prior to the physiological regulation of cGMP channels, likely results from altered energy homeostasis and abnormal metabolic and calcium signaling. Our findings provide a plausible framework for the etiology of retinal degeneration by incorporating unique photoreceptor ‘omics’ and physiology in concert with our observations from the *rd1* mutant retina. In addition, our studies are consistent with the mechanistic underpinning of neurodegenerative diseases ^62,63^ and should have broad implications for deciphering molecular and cellular drivers of disease pathology. Discovery of metabolic and signaling pathways that are altered in anticipation of disease sets the foundation for drug discovery and interventions at an early stage, thereby improving outcomes in clinical management.

## METHODS

### Animal models

Mice were housed in 12-hour light-dark housing conditions and were cared for following recommendations of the Guide for the Care and Use of Laboratory Animals, Institute of Laboratory Animal Resources, the Public Health Service Policy on Humane Care and Use of Laboratory Animals. All mouse protocols have been approved by the Animal Care and Use Committee of the National Eye Institute (ASP#650). The *Pde6b^rd1/rd1^; TgNrlp-EGFP* mice (referred to as *rd1*) were generated by crossing *Pde6b^rd1/rd1^* mice to *TgNrlp-EGFP* mice until homozygous and maintained as sibling crosses. *TgNrlp-EGFP* mice served as wild type (*WT*) control and were maintained on C57BL/6J backgrounds with >10 backcrosses. Both male and female mice were used in this study.

### Retinal histology

After enucleation, eyes were fixed with 4% glutaraldehyde for 30 min at room temperature and then with 4% paraformaldehyde overnight at 4°C. Eyes were then embedded in methacrylate and sectioned at 1-micron thickness. Representative sections from the superior central retina were stained with standard hematoxylin and eosin (H&E) staining protocol. Pictures of retinal section were taken at 40X.

### Transcriptomic profiling

#### Flow sorting of rod photoreceptors

Rod photoreceptors from *Pde6b^rd1/rd1^; TgNrlp-GFP* (*rd1*) and *TgNrlp-GFP* (*WT*) mice were isolated as previously described ^33^. In brief, dissected retinas were treated with papain (Worthington Biochemical, NJ, USA) and dissociated singe cells were collected by centrifugation at 800 x g for 5 min at 4°C. Cell pellets were resuspended in ice cold PBS. GFP-positive photoreceptor cells were isolated using FACS Aria II (Becton Dickinson, CA, USA) with a stringent precision setting which maximized the purity of sorted cells.

#### RNA sequencing

Rod photoreceptor cells were lysed with TRIzol LS (Invitrogen), and the RNA was isolated following the manufacturer’s instructions. RNAseq data were generated using TruSeq Stranded mRNA Sample Prep Kit (Illumina), as previously described ^64^, and 125 base pair-end reads were generated on HiSeq 2500 platform (Illumina).

#### Differential RNA expression and downstream analysis

Unprocessed fastq files from the sequencer were quality checked, trimmed and aligned to the *Mus musculus* Ensembl annotation (v98) using a robust pipeline as described before ^64^. Briefly, the count matrix was normalized, and batch corrected to generate abundance summary in counts per million using the Bioconductor packages *edgeR* and *limma*. Additionally, to remove artefacts we set an expression filter that required genes to have greater than 5 CPM in minimum two replicates of any sample before qualifying for differential expression analysis. Statistical tests for differential expression between age-matched samples of *rd1* and *WT* were performed with *voom* transformed counts fit to a linear model using functions in *limma*. Finally, a cutoff of 10% FDR and 1.5-fold change difference was decided to group genes as differentially expressed genes (DEG). Clusters of DEG were determined from a k-means algorithm after setting a value of k equals 10. Functional enrichment analysis of DEG groups was performed with *gProfileR* and custom scripts where KEGG pathways were ordered by pathway impact, as defined by the ratio between overlap and pathway size. Gene sets for volcano plots were obtained from gProfileR (Reactome), Mouse Genomics Institute (Gene Ontology) and Mitocarta.

### Proteomic profiling

#### Retinal tissue lysate processing

Retinal tissues from *WT* and *rd1* mice at P6 and P10 were subject for whole proteomic analysis, performed by core facility at National Heart Lung and Blood Institute. Retinas from 9 non-littermate mice of each sample group were frozen and thawed after dissection, divided in replicates of 6 retinas for each strain and age (n=3 for each group,12 samples total), and resuspended in 100 μl of lysis buffer containing 6 M urea, 2 M thiourea and 4% CHAPS (Thermo Fisher Scientific). Each combined replicate of samples was loaded onto a QIAshredder spin column (QIAGEN) placed in a 2 ml collection tube and spun for 2 minutes at maximum speed in a microcentrifuge to aid in tissue disruption. The flow-through for each sample was collected and protein was quantified using the Pierce detergent compatible Bradford assay kit (Thermo Fisher Scientific).

A volume corresponding to 100 μg of protein from each replicate, as well as two normalization loading controls generated by pooling 5.55 μg of each replicate were taken to 100 μl by adding the necessary volume of lysis buffer. Each resulting sample was reduced by mixing with 5 μl of the 200 mM DTT and incubating at room temperature for 1 hour, and then alkylated by adding 5 μl of 375 mM iodoacetamide and incubating for 30 minutes protected from light at room temperature. Protein in each sample was precipitated by adding 600 μl of pre-chilled acetone and storing overnight at −20°C. Samples were then centrifuged at 8000 × g for 10 min at 4°C, and acetone was carefully decanted without disturbing the white pellet, which was allowed to dry for 2-3 minutes. Each protein pellet was resuspended in 100 μl of a buffer containing 100 mM triethylammonium bicarbonate (TEAB) and 0.1% of Progenta anionic acid-labile surfactant I (AALSI, Protea Biosciences). Each sample was proteolyzed by adding 10 μl of 1.25 μg/μl sequencing grade modified trypsin (Promega) dissolved in 100 mM TEAB and incubated overnight at 37°C. Each sample was labeled with a different TMT label reagent (Thermo Fisher Scientific) by adding the contents of each label tube after dissolving with 41 μl of acetonitrile. The reaction was allowed to proceed for 1 hour at room temperature and was quenched by adding 8 μl of 5% hydroxylamine to each sample and incubating for 15 minutes.

All seven samples from each strain (6 replicates plus a normalization loading control) were combined in a new microcentrifuge tube, which was dried under vacuum until all acetonitrile was removed. Contaminating CHAPS was removed using 2 ml Pierce detergent removal columns (thermo Fisher Scientific) and residual AALSI was cleaved by adding 30 μl of 10% trifluoroacetic acid (TFA). The combined samples were desalted using an Oasis HLB column (Waters) and dried under vacuum.

#### Offline HPLC peptide fractionation

High pH reversed-phase liquid chromatography was performed on an offline Agilent 1200 series HPLC. Approximately 1 mg of desalted peptides were resuspended in 0.1 ml of 10 mM triethyl ammonium bicarbonate with 2% (v/v) acetonitrile. Peptides were loaded onto an Xbridge C_18_ HPLC column (Waters; 2.1 mm inner diameter x 100 mm, 5 μm particle size), and profiled with a linear gradient of 5–35 % buffer B (90% acetonitrile, 10 mM triethyl ammonium bicarbonate) over 60 minutes, at a flowrate of 0.25 ml/min. The chromatographic performance was monitored by sampling the eluate with a diode array detector (1200 series HPLC, Agilent) scanning between wavelengths of 200 and 400 nm. Fractions were collected at 1 minute intervals followed by fraction concatenation ^65^. Fifteen concatenated fractions were dried and resuspended in 0.01% formic acid, 2% acetonitrile. Approximately 500 ng of peptide mixture was loaded per liquid chromatography-mass spectrometry run.

#### Mass Spectrometry (MS)

All fractions were analyzed on an Ultimate 3000-nLC coupled to an Orbitrap Fusion Lumos Tribrid instrument (Thermo Fisher Scientific) equipped with a nanoelectrospray source. Peptides were separated on an EASY-Spray C_18_ column (75 μm x 50 cm inner diameter, 2 μm particle size and 100 Å pore size, Thermo Fisher Scientific). Peptide fractions were placed in an autosampler and separation was achieved by 90 minutes gradient from 4-35% buffer B (100% ACN and 0.1% formic acid) at a flow rate of 300 nL/min. An electrospray voltage of 1.9 kV was applied to the eluent via the EASY-Spray column electrode. The Lumos was operated in positive ion data-dependent mode, using Synchronous Precursor Selection (SPS-MS3) ^66^.

Full scan MS1 was performed in the Orbitrap with a precursor selection range of 375 to 1,275 m/z at 1.2 x 10^5^ normal resolution. The AGC target and maximum accumulation time settings were set to 4 x 10^5^ and 50 ms, respectively. MS2 was triggered by selecting the most intense precursor ions above an intensity threshold of 5 x 10^3^ for collision induced dissociation (CID)-MS^2^ fragmentation with an AGC target and maximum accumulation time settings of 1 x 10^4^ and 60 ms, respectively. Mass filtering was performed by the quadrupole with 0.7 m/z transmission window, followed by CID fragmentation in the linear ion trap with 35% normalized collision energy in turbo scan mode and parallelizable time option was selected. SPS was applied to co-select 10 fragment ions for HCD-MS3 analysis. SPS ions were all selected within the 400–1,200 m/z range and were set to preclude selection of the precursor ion and TMTC ion series. The AGC target and maximum accumulation time were set to 1 x 10^5^ and 125 ms (respectively) and parallelizable time option was selected. Co-selected precursors for SPS-MS^3^ underwent HCD fragmentation with 65% normalized collision energy and were analyzed in the Orbitrap with nominal resolution of 5 x 10^4^. The number of SPS-MS^3^ spectra acquired between full scans was restricted to a duty cycle of 3 s.

#### Mass spectrometry data processing

Raw data files were processed using Proteome Discoverer (v2.4, Thermo Fisher Scientific), using both Mascot (v2.6.2, Matrix Science) and Sequest HT (Thermo Fisher Scientific) search algorithms. All the peak lists were searched against the UniProtKB/Swiss-Prot protein database released 2020_02 with *Mus musculus* taxonomy (17,010 sequences) and concatenated with reversed copies of all sequences. The following search parameters were set as static modifications; carbamidomethylation of cysteine, TMT 10-plex modification of lysine and peptide N-terminus. The variable modifications were set as; oxidation of methionine, deamidation of aspartamine and glutamine. For SPS-MS3 the precursor and fragment ion tolerances of 12 ppm and 0.5 Da were applied, respectively. Up to two-missed tryptic cleavages were permitted. Percolator (v3.02.1, University of Washington) algorithm was used to calculate the false discovery rate (FDR) of peptide spectrum matches, set to q-value 0.05 ^67,68^. TMT 10-plex quantification was also performed by Proteome Discoverer v.2.4 by calculating the sum of centroided ions within 20 ppm window around the expected m/z for each of the 10 TMT reporter ions. Spectra with at least 50% of SPS masses matching to the identified peptide are considered as quantifiable PSMs. Quantification was performed at the MS^3^ level where the median of all quantifiable PSMs for each protein group was used for protein ratios.

#### Differential expression analysis of the proteome

Proteins identified from the MS experiment were filtered to retain only those with at least one unique peptide and that have minimum two peptide spectral matches from either of the two algorithms, MASCOT or Sequest. Further, to classify proteins as differentially expressed, a threshold of 1.2-fold change in both directions of over- and under-expression was decided for age-matched comparisons between *rd1* and *WT*.

#### Pathway enrichment analysis of the proteome

GSEA for the proteomic dataset was performed using *fgsea* with ranked fold change values and Reactome and Gene Ontology annotations downloaded from the gProfileR webserver. *Fgsea* significant pathways (*p*<0.05) were further collapsed to address redundancy. Ridge plots were generated using custom scripts and functions in the *tidyverse* and *ggridges* packages. Additionally, a list of OXPHOS proteins and electron transport chain complexes were curated from literature for use in various analyses.

### Metabolomic profiling

#### Retinal metabolome profiling

Targeted quantitative analysis was performed on retinal samples from *WT* and *rd1* mice of P6 and P10 to detect cationic and anionic metabolites, using capillary electrophoresis mass spectrometry (CE-TOFMS and CE-QqQMS) in cation and anion analysis modes, respectively, with internal standard (Human Metabolome Technologies). Retinal tissue samples (30-45 mg) were mixed with 50% (v/v) acetonitrile and homogenized and the supernatant obtained after brief centrifugation was filtrated through 5 kDa cut off filter to remove macromolecules. Filtrate was centrifugally concentrated and resuspended in water for mass spectrometry measurement. Metabolite concentrations were calculated by normalizing to internal standard and quantity of sample used. Primary analysis of metabolite detection and abundance estimates were performed by Human Metabolome Technologies, Boston MA.

#### Retinal metabolome analysis

For analysis, metabolites were linked to their corresponding KEGG pathways and visualized in a network diagram created with *igraph* (R package) and Cytoscape. To classify differential abundance of metabolites we used two sets of thresholds: a) significance differential abundance with *p*<0.05 from pairwise tests of age-matched samples at P6 and P10; and b) metabolites with fold change of minimum 1.2 times in either over or under abundance in *rd1* versus *WT* comparisons.

### In vitro and ex vivo analysis of mitochondrial activity

#### Complex I activity assay

The protocol for complex I activity assay was adopted from previous publications ^69^. In brief, dissected retinas were homogenized in SETH buffer containing 0.25 M sucrose, 2 mM EDTA and 10 mM Tris pH7.4. The lysate was first spun at 600 x g for 10 min at 4°C followed by a second spin of supernatant at 14000 x g for 10 min at 4°C. Mitochondria pellet was resuspended in 10 mM Tris pH7.4. Complex I oxidizes NADH and produces an electron. The produced electron is then used to reduce an artificial substrate decylubiquinone, which subsequently passes the electron to the terminal electron receptor DCIP that is blue in color. Reduction of DCIP concentration can be followed spectrophotometrically at 600nm. Rotenone is added to terminate the reaction and reveal any residue non-complex I related reduction of DCIP.

#### Mitochondrial respiration and oxygen consumption

OCR of retinal punches were measured using Seahorse XF24 Bioanalyzer with XF24 Islet Fluxpaks (Agilent) following our published protocol ^9^. In brief, eyes were enucleated and placed in ice-cold PBS for dissection. Cornea and lens were removed, and the retinal cup was gently separated away from the scleral layer. Three to four of 1 mm diameter punches equidistant from the optic nerve head were taken from each retina for the measurement. Ames’ buffer (Sigma) containing 120 mM NaCl, 10 mM HEPES, pH 7.4 was used in the assay. After steady basal OCR was established, mitochondrial uncoupler (2-fluorophenyl)(6-[(2-fluorophenyl)amino](1,2,5-oxadiazolo[3,4-e]pyrazin-5-yl))amine (BAM15; Timtec, Newark, DE, USA) was added at a final concentration of 5 μM to uncouple the electrochemical gradient of protons, thereby uncoupling ATP production from oxygen consumption and causing mitochondrial OXPHOS to run at full capacity. The complex I inhibitor rotenone was used at a final concentration of 1 μM to inhibit the entire electron transport chain and thereby revealing the residual non-mitochondrial oxygen consumption.

Raw OCR values were computed by the Seahorse KSV algorithm, per manufacturer’s recommendations, and non-mitochondrial oxygen consumption, the residual readings obtained after addition of rotenone, was subtracted from all points. Basal mitochondrial OCR (OCRbasal) was taken at 36 min, the last measuring point before BAM15 addition, while the highest value from uncoupled points was used for maximal OCR (OCR_max_). Mitochondrial reserve respiratory capacity (MRC) was calculated as the percentage difference between maximal uncontrolled oxygen consumption rate OCR_max_ and the initial 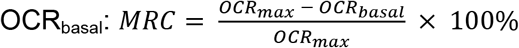. A range of 4 to 8 non-littermate animals were included in each age group of *WT* and *rd1* mice, and data from 12~29 retinal punches were included for analysis in each age/genotype group. Group means ± SEMs are plotted for each age group and strain.

#### Transmission Electron Microscopy

The mitochondrial ultrastructure was evaluated by transmission electron microscopy (TEM) following standard preparation procedures ^70^. Eyes were enucleated and immersion fixed in 1% paraformaldehyde, 2.5% glutaraldehyde in PBS (pH 7.4), followed by treatment with 1% osmium tetroxide. Fixed samples were dehydrated through a series of ethanol gradient and in propylene oxide, and then embedded in epoxy resin. The retina samples were sectioned at 80-90 nm, mounted on copper grids, and doubly stained with uranyl acetate and lead citrate. Sections were imaged at the NEI Histology Core with JEOL JEM-1010 TEM. Photographs were taken for the inner segment region of the photoreceptors at 10,000x and 30,000x magnifications.

#### qPCR and Mit/Nu DNA ratio

The Mit/Nu DNA ratio were determined on P2 to P10 retinal samples from *WT* and *rd1* mice using a published protocol ^37^. Whole genomic DNA was isolated from retinal tissues and quantitative PCR on the genomic DNA abondance of mitochondria-encoded *Nd1* and nucleus-encoded *Hk2* was performed to evaluate copy number of mitochondrial and nuclear genome respectively. The mtDNA to nuDNA ratio was calculated by a ΔΔCt method.

### Quantification and statistical analysis

Unless specified otherwise, all statistical analyses and data visualizations were done using the R statistical platform. Differences from *WT* were regarded as significant if p < 0.05.

## ACKNOWLEDGMENTS

We are grateful to Yide Mi, Ashley Yedlicka and Megan Kopera for mouse colony management; Julie Laux, Jessica Albrecht and Rafael Villasmil of the NEI flow cytometry core for their assistance; and Thad Whitaker, Maria Campos, Mones Abu-Asab and the NEI histology core for histology and TEM. This research was supported by Intramural Research Program of the NEI (ZIAEY000450 and ZIAEY000546) and utilized the high-performance computational capabilities of the Biowulf Linux cluster at NIH (http://biowulf.nih.gov).

## Author Contributions

Overall Conceptualization, K.J., J.W.K., A.K.M., R.C. and A.S.; Methodology and Investigation, K.J., Y.A., J.G., J.W.K., A.B., J.N.; Omics Data Generation and Analysis, A.K.M., A.A., L.G., M.J.B., R.C.; Data Curation, A.K.M.; Resources, K.J., D.A.F., R.B., R.C., Writing – Original Draft, K.J., A.K.M., A.S.; Writing – Review & Editing, all authors; Supervision, R.C., A.S.; Funding Acquisition and Project Administration, A.S.

## Materials and Correspondence

Correspondence and material requests should be addressed to Anand Swaroop, Ph.D., Neurobiology-Neurodegeneration & Repair Laboratory, National Eye Institute, National Institutes of Health, MSC0610, 6 Center Drive, Bethesda, MD, USA. Phone: 301.435.5754; Fax: 301.480.9917;

