## Supplementary Figures and Legends for "Early mitochondrial stress and metabolic imbalance lead to photoreceptor cell death in retinal degeneration"

**Early mitochondrial defects and metabolic imbalance precede the onset of retinal neurodegeneration**

**CONTENTS**

Supplementary figures

Supplementary data legends

### Supplementary Figure 1

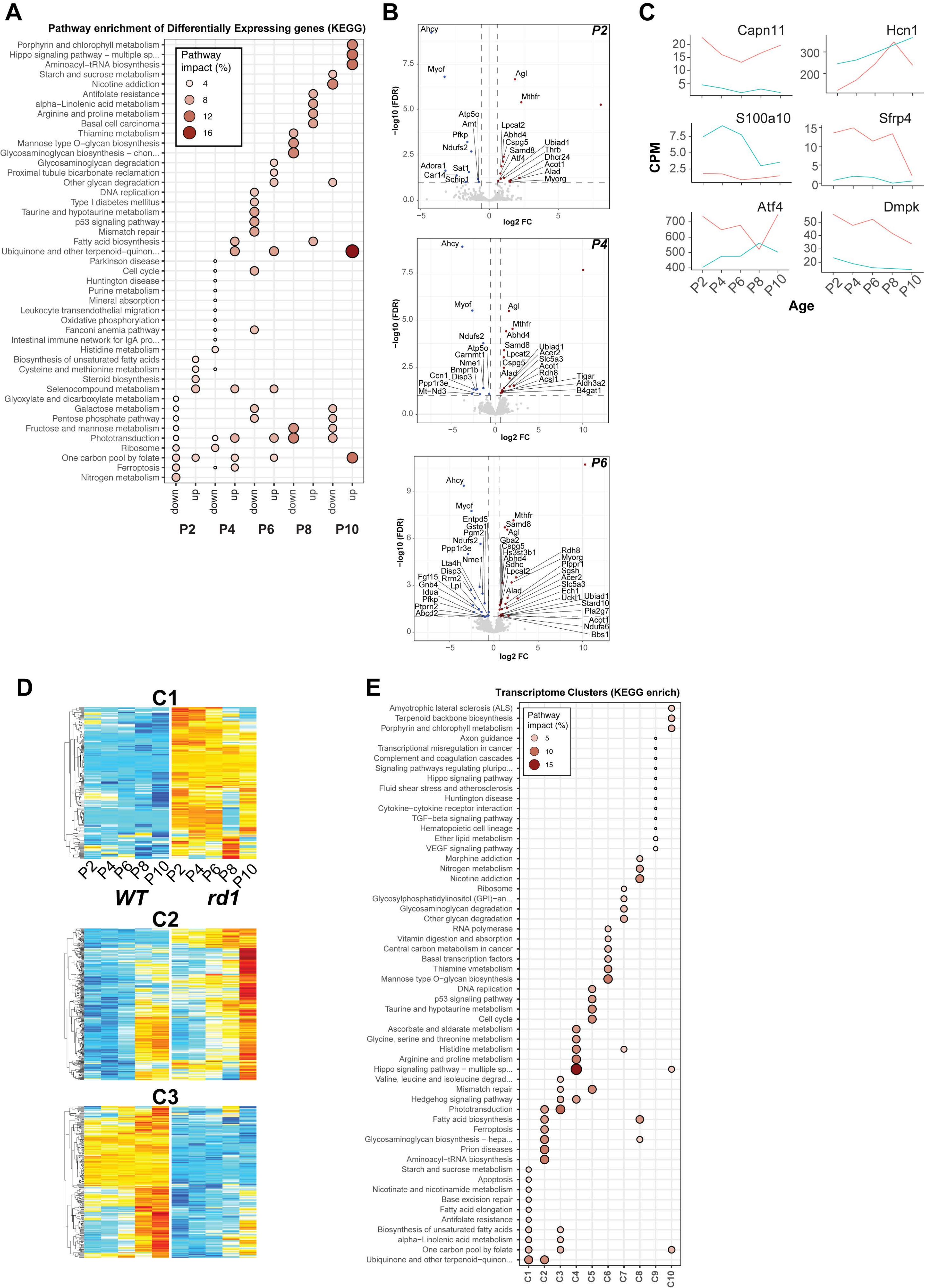

**Supplementary Figure 1. Differential gene expression in *rd1* rod photoreceptors at pre-degeneration stages.**

A. Enrichment plot of top 10 KEGG pathways that are most impacted by differential gene expression at pre-degeneration stages in *rd1* rods.
B. Volcano plots of metabolic genes (as annotated in Reactome) at P2, P4, and P6. Colors represent significance and direction of differential expression (*rd1* vs *WT*): red – significantly over-expressed; blue – significantly under-expressed; and, grey – not significant.
C. Calcium-related and signaling genes show divergent expression in *rd1* rod photoreceptors at neonatal stages. Red and blue lines denote gene expression in *rd1* and *WT* rod photoreceptors, respectively.
D. Heatmaps of selected clusters - C1, C2 and C3, showing atypical gene expression trends between *rd1* and *WT* rods. Log CPM (lcpm) values are row scaled to z-scores for plotting. Color scale bar is same as in Figure 1D.
E. Pathway annotation (KEGG) of clusters of differentially expressed genes between rod photoreceptors from *rd1* and *WT* retina. The dot plot shows the top 10 most impacted pathways per cluster.

### Supplementary Figure 2

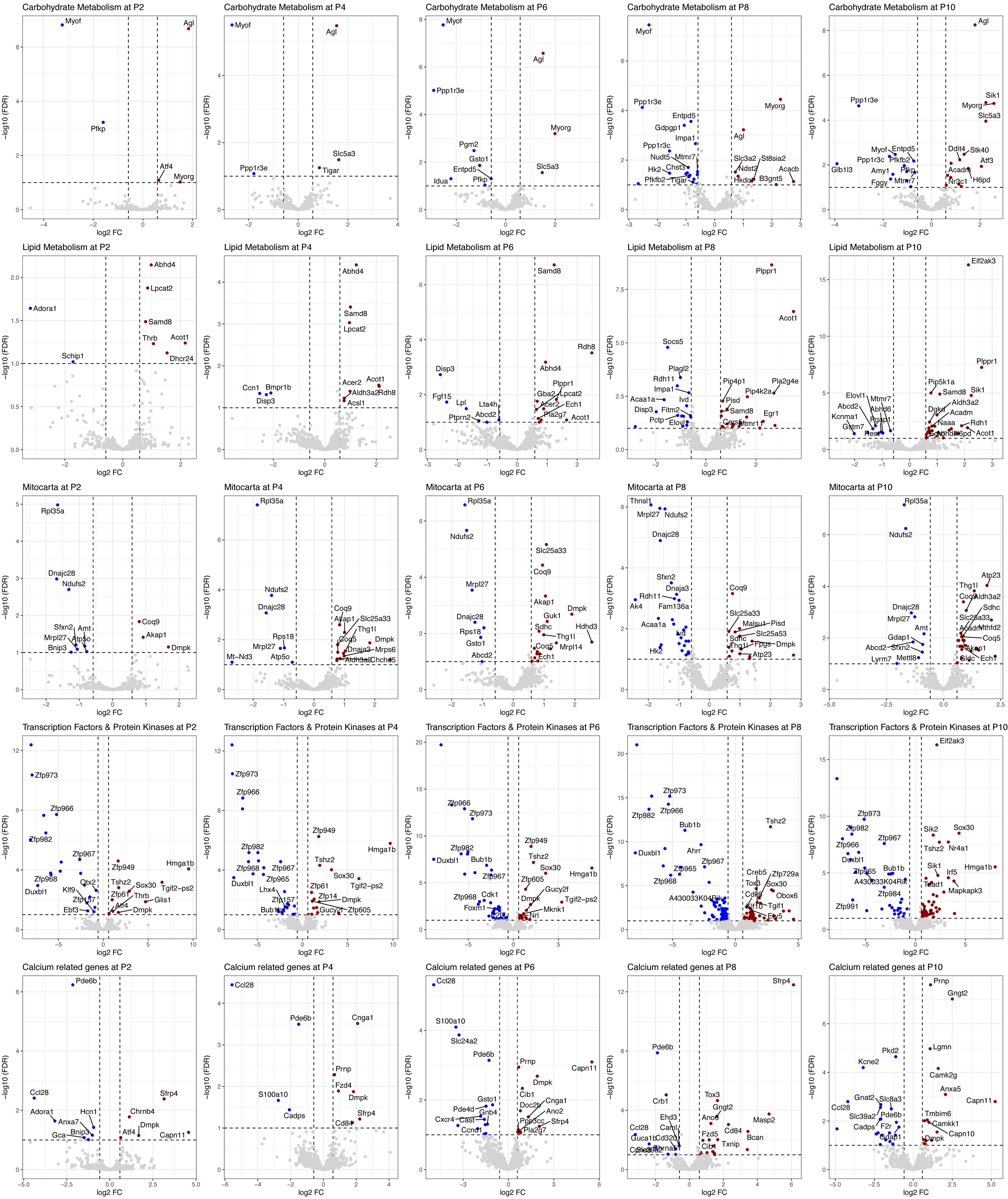

**Supplementary Figure 2. Transcriptional deregulation in selected pathways before rod photoreceptor degeneration in the *rd1* retina.** Volcano plots summarizing differential expression of selected pathways – Carbohydrate metabolism, lipid metabolism, mitocarta, transcription factors and protein kinases, and calcium related genes, at P2, P4, P6, P8 and P10, in rod photoreceptors from *rd1* retina before degeneration. Colors represent significance and direction of differential expression (*rd1* vs *WT*): red – significantly over-expressed; blue – significantly under-expressed; and, grey – not significant.

### Supplementary Figure 3

**A**

**Top overexpressed proteins (P6)**

| Symbol | Description | FC |
|---|---|---|
| CAR8 | Carbonic anhydrase 8 | 100 |
| HDDC3 | HD domain containing 3 | 14.322 |
| CD99L2 | CD99 antigen-like 2 | 12.285 |
| ACOT1 | Acyl-CoA thioesterase 1 | 5.19 |
| SUPV3L1 | Suppressor of var1, 3-like 1 | 4.98 |

**Top underexpressed proteins (P6)**

| Symbol | Description | FC |
|---|---|---|
| AMY2A5 | Amylase 2a5 | 0.055 |
| CALM1 | Calmodulin 1 | 0.127 |
| PDE6B | Phosphodiesterase 6B, cGMP-specific, rod, beta | 0.138 |
| AQP4 | Aquaporin 4 | 0.219 |
| CRYAA | Crystallin, alpha A | 0.228 |

**Top overexpressed proteins (P10)**

| Symbol | Description | FC |
|---|---|---|
| HDDC3 | HD domain containing 3 | 15.838 |
| CD99L2 | CD99 antigen-like 2 | 6.042 |
| ACOT1 | Acyl-CoA thioesterase 1 | 4.416 |
| SUPV3L1 | Suppressor of var1, 3-like 1 (S. cerevisiae) | 3.137 |
| RNF146 | Ring finger protein 146 | 2.999 |

**Top underexpressed proteins (P10)**

| Symbol | Description | FC |
|---|---|---|
| PDE6B | Phosphodiesterase 6B, cGMP-specific, rod, beta | 0.041 |
| AMY2A5 | Amylase 2a5 | 0.089 |
| PDE6A | Phosphodiesterase 6A, cGMP-specific, rod, alpha | 0.135 |
| CRYGS | Crystallin, gamma S | 0.284 |
| RAB33B | RAB33B, member RAS oncogene family | 0.293 |

**B**

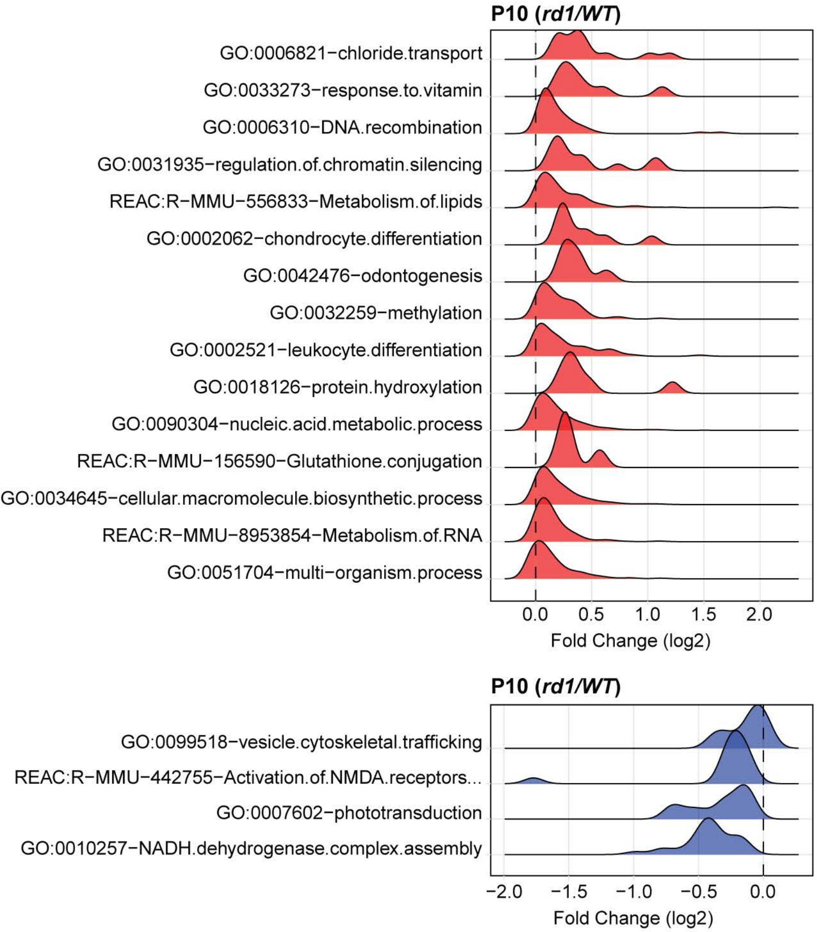


**C**

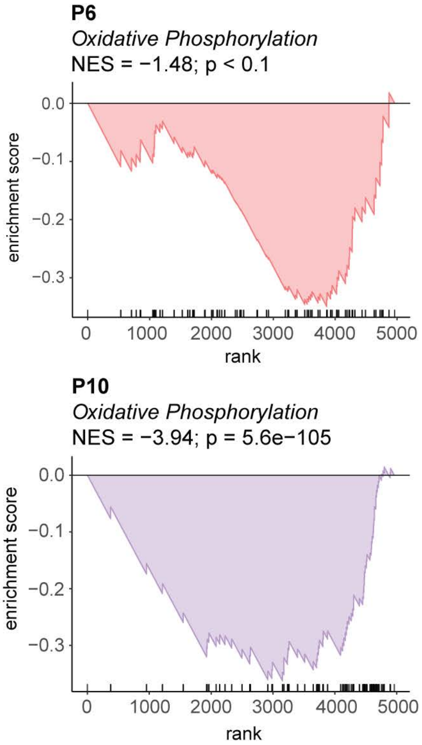


**D**

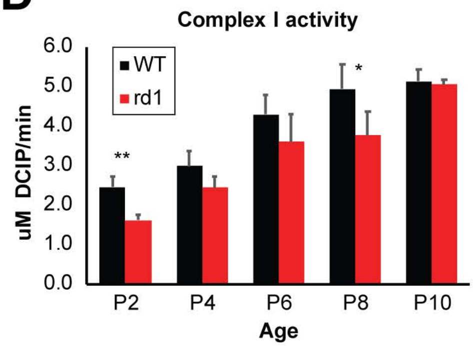

**Supplementary Figure 3. Proteomic changes in the *rd1* retina and decreased activity of mitochondrial NADH-dehydrogenase complex (Complex I).**

A. A list of the top five over- and under-expressing proteins at P6 and P10 in the *rd1* retina.

B. Ridge-plots of significantly differential pathways at P10 from gene set enrichment analysis. The top (red fill) and bottom (blue fill) panels represent over- and under-enriched pathways, respectively. Each ridge shows distribution of fold change of leading-edge proteins of the respective significant pathway.

C. Enrichment curve from *fgsea* analysis for a curated list of OXPHOS proteins in *rd*1 vs *WT* retina at P6 and P10, respectively.

D. Reduced Complex I activity in the *rd1* retina before photoreceptor degeneration. Complex I activity assayed in mitochondrial enriched *WT* and *rd1* retinal lysates, n=3 or 4 for each group. Data are represented as mean ± SEM. Student t test, * $p$<0.05, ** $p$<0.01.

### Supplementary Figure 4

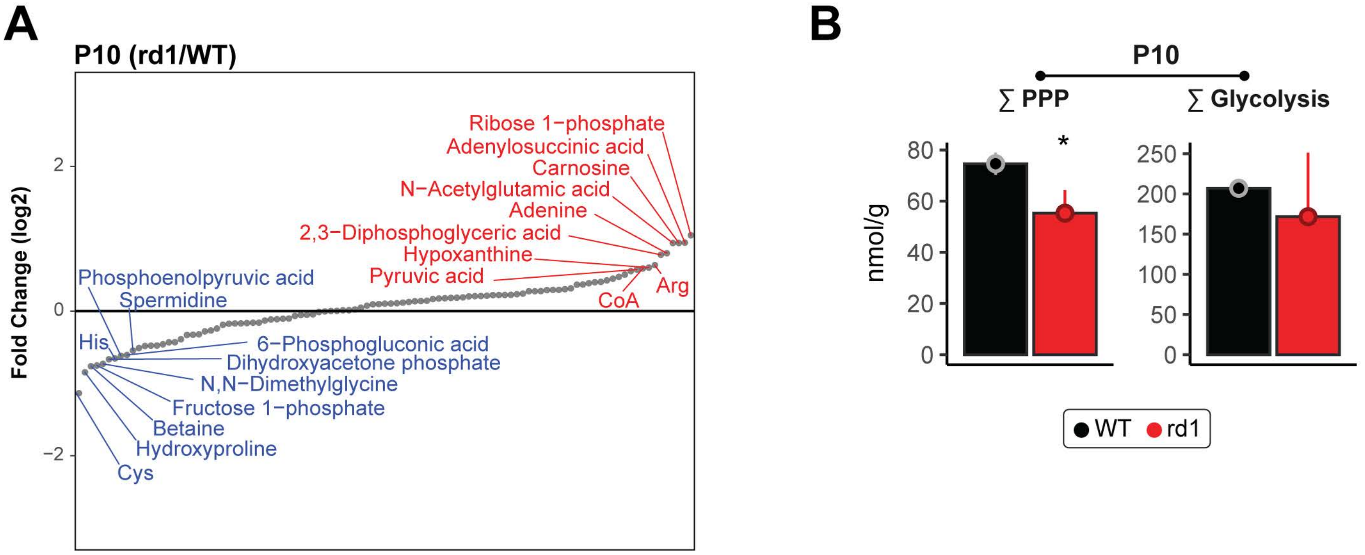

**Supplementary Figure 4. Aberrant flux in central carbon metabolism continues at P10.**

A. Plot showing metabolite abundance profiles of the *rd1* retina relative to *WT*, at P10. Red labels denote top abundant metabolites in the *rd1* retina, while blue labels denote most depleted metabolites.

B. Reduced glycolytic and pentose phosphate pathway flux as seen by cumulative metabolite abundance at P10. Color and significance features are same as Figure 4C.

### Supplementary Figure 5

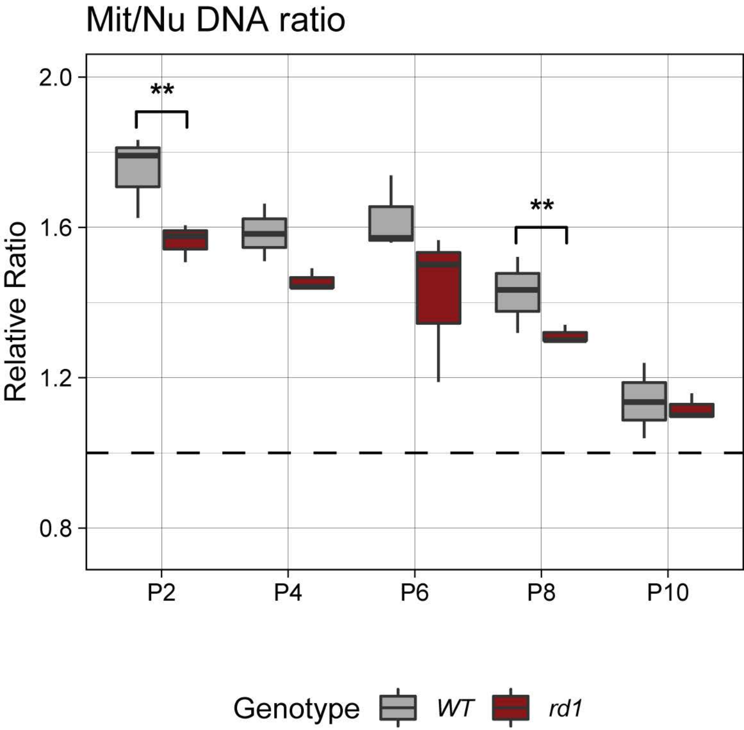

**Supplementary Figure 5. Lower Mit/Nu DNA ratio in *rd1* retinas.** Mit/Nu DNA ratio was measured in *WT* and *rd1* retinas from P2 to P10. Student t test, ** $p<0.01$.

### Supplementary Figure 6

**TEM of *rd1* retina showing ONL, Outer segment and RPE**

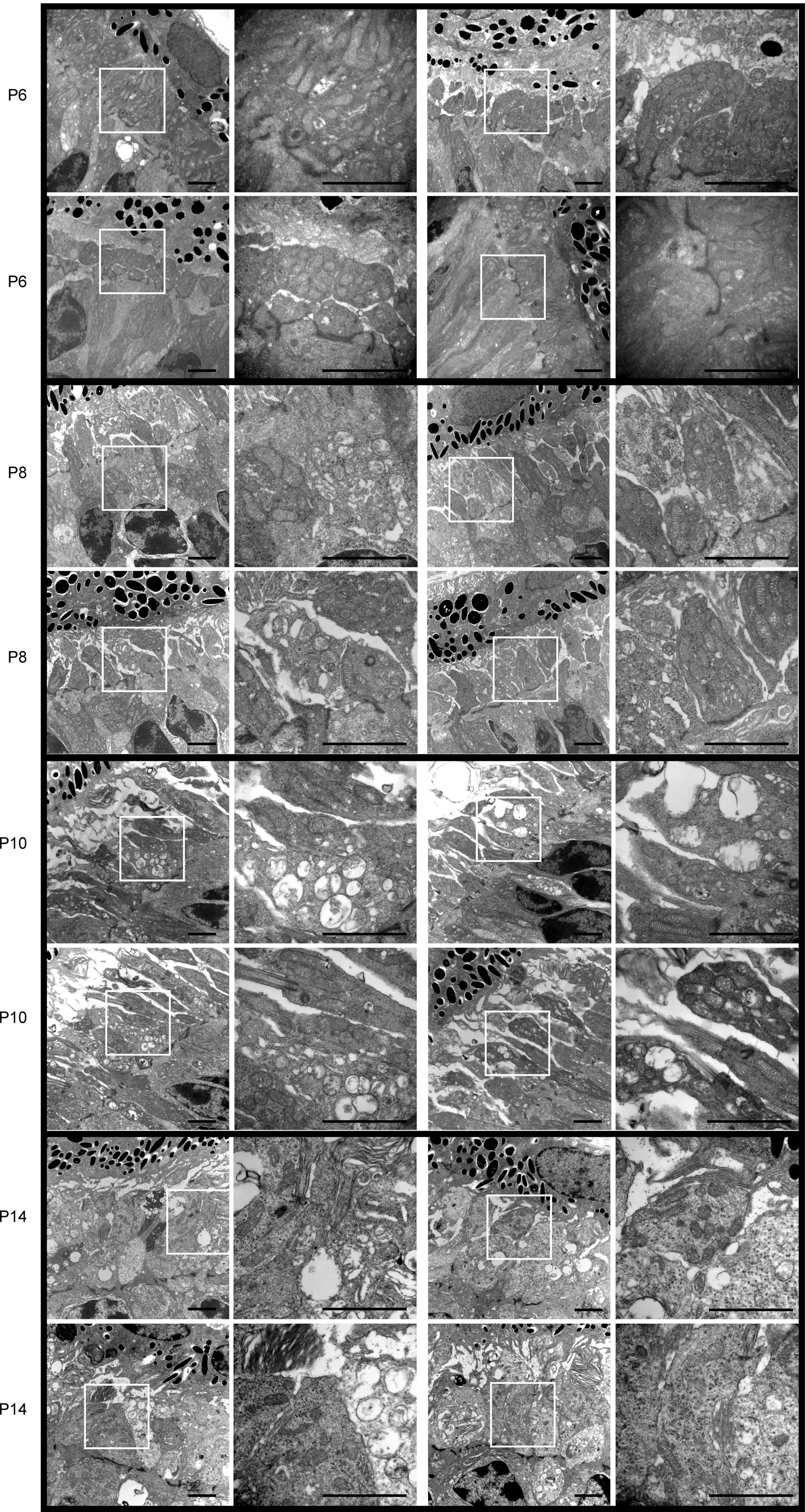

**Supplementary Figure 6. TEM ultrastructure assessment of photoreceptor inner segments and mitochondria in *rd1* photoreceptors from P6 to P14.** Photographs were taken at 10,000X and 30,000X. White square box indicates zoomed in area. Scale bar = 2 mm.

#### SUPPLEMENTARY DATA LEGENDS

**Supplementary Data 1. Gene expression quantification of flow sorted rod photoreceptors from *WT* and *rd1* retina.** RNAseq summary in counts per million (CPM) and statistical test results for significantly differentially expressed genes between flow sorted rod photoreceptors from *WT* and *rd1* retina.

**Supplementary Data 2. Proteomic quantitation and differential analysis of *WT* and *rd1* retina.** Peptide detection summary and abundance ratio for retinal proteins detected in the mass spectrometry experiment.

**Supplementary Data 3. Retinal metabolome landscape in *WT* and *rd1*.** Summary of comparative analysis of metabolomic profiles from *WT* and *rd1* retina.
